# The Imaging and Molecular Annotation of Xenografts and Tumours (IMAXT) High Throughput Data and Analysis Infrastructure

**DOI:** 10.1101/2021.06.22.448403

**Authors:** Eduardo A. González-Solares, Ali Dariush, Carlos González-Fernández, Aybüke Küpcü Yoldaş, Mohammad Al Sa’d, Neil Millar, Tristan Whitmarsh, Nicholas Chornay, Ilaria Falciatori, Atefeh Fatemi, Daniel Goodwin, Laura Kuett, Claire M. Mulvey, Marta Páez Ribes, Fatime Qosaj, Andrew Roth, Ignacio Vázquez-García, Spencer S. Watson, Jonas Windhager, Samuel Aparicio, Bernd Bodenmiller, Ed Boyden, Carlos Caldas, Owen Harris, Sohrab P. Shah, Simon Tavaré, CRUK IMAXT Grand Challenge Team, Dario Bressan, Gregory J. Hannon, Nicholas A. Walton

## Abstract

With the aim of producing a 3D representation of tumours, IMAXT uses a large variety of modalities in order to acquire tumour samples and produce a map of every cell in the tumour and its host environment. With the large volume and variety of data produced in the project we develop automatic data workflows and analysis pipelines and introduce a research methodology where scientists connect to a cloud environment to perform analysis close to where data are located instead of bringing data to their local computers. Here we present the data and analysis infrastructure, discuss the unique computational challenges and describe the analysis chains developed and deployed to generate molecularly annotated tumour models. Registration is achieved by use of a novel technique involving spherical fiducial marks that are visible in all imaging modalities used within IMAXT.

## Main

Single cell analysis providing a detailed genomic and proteomic breakdown of tissues is now well established. Hitherto, spatial information has been lost. Recognising the importance of understanding the detailed environments of tumours, the Cancer Grand Challenge identified a key challenge to map the molecular and cellular tumour microenvironment (https://cancergrandchallenges.org/challenges/3d-tumour-mapping) in order to define new targets for therapy and prognosis. The Imaging and Molecular Annotation of Xenografts and Tumours (IMAXT) project is adopting an integrated approach to study tumours and their environment building a 3D representation that can be explored using virtual reality and show every single, fully annotated cell type, in the tumour and surroundings.

To do this the project uses a large variety of technologies and instrumental modalities gathering multi-disciplinary expertise from many international groups including sequencing, molecular biology, statistics, medicine, astronomy and virtual reality experts.

Serial Two-Photon Tomography^1^ (STPT) is the fastest and most high-throughput of the IMAXT data acquisition modalities, and the only one capable of processing sample numbers in the range of hundreds of full-size (centimetre-level) tumours, or thousands of biopsies, providing full 3D models at single cell resolution. It also serves as the starting point and sectioning step for our deeper analysis pipelines, including Imaging Mass Cytometry^2^ (IMC), and, in the near future, Expansion Sequencing^3^ (ExSeq) and Multiplexed error-robust fluorescence in situ hybridization^4^ (MERFISH) (which are typically performed on frozen sections). These are complemented by single cell RNA and DNA sequencing.

Figure 1 shows the data acquisition and analysis workflow that has been implemented. A tumour sample is collected (via biopsy or resection from a mouse implant) and embedded in agarose in order to maintain the sample integrity and allow for sectioning. At the same time fluorescent spherical agarose beads about 90 µm in diameter are inserted in the cube in the areas not covered by the sample. These beads will be especially useful during image registration (see Methods). The size of the final block is around 1 cm^3^. Slices as thin as 15 µm are sectioned using a vibratome of the TissueCyte 2000 instrument (TissueVision Inc, Newton, MA, USA) and imaged through a variety of instruments. STPT performs two-photon fluorescence imaging in four channels a few microns below the surface of the cube. The cube is then cut and each slice imaged with fluorescence scanning (Zeiss Axioscan slide scanner) in several channels. The slice is then acquired with IMC which provides information on up to 40 individual metal-conjugated antibodies. Slices are registered using the beads inserted in the agarose cube and all data is resampled to the STPT reference frame. Together with sequencing data all data is then federated to make an annotated 3D model with each cell having tens to hundreds of descriptors.

**Figure 1:**
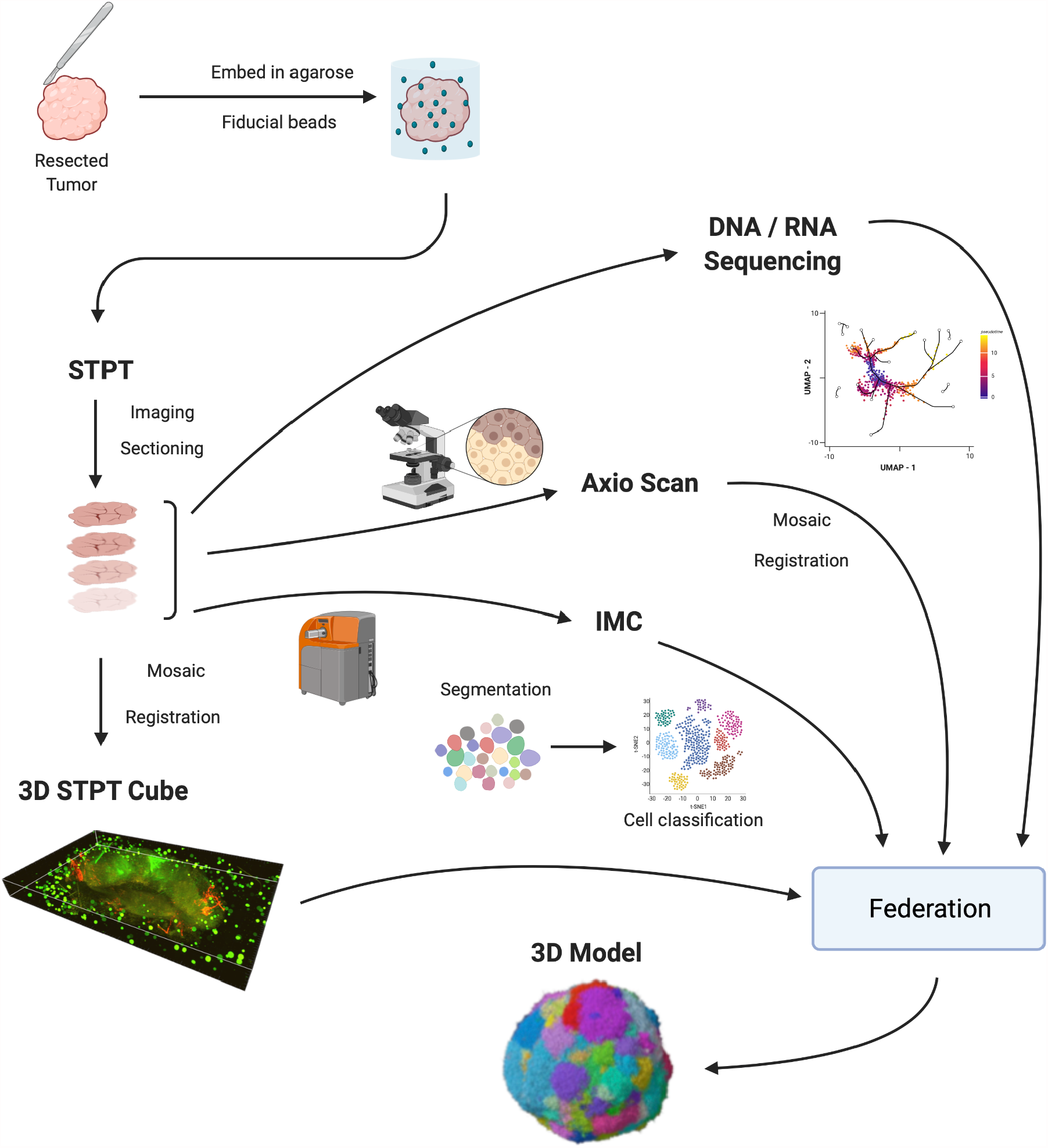
IMAXT pipeline. Once a tumour has been extracted it is embedded in an agarose cube together with spherical beads. The sample is then analysed in the STPT instrument where it is imaged and cut into thin slices and a multichannel 3D data cube is produced. The slices are then imaged with an Axioscan fluorescence microscope and an IMC mass cytometer. The spherical beads are used for alignment of slices within each sample and for registration between all samples. All imaging is resampled to the STPT reference. All data, including sequencing, is then federated to build an annotated 3D model.

Many of these technologies produce large quantities of data that need to be processed before any scientific analysis can be performed. As the technologies mature, rates of data production also increase. As an example STPT alone generates between 2 to 3TB of imaging data per day for a single sample. Analysis of these large volumes of data, that are available at high data rates, calls for automatic processing pipelines that can optimally run with high a level of parallelism and produce results in a timely fashion.

IMAXT also presents a number of unique computational challenges, where approaches developed for data handling and image analysis of multi-wavelength survey data in astronomy (e.g.^5^ for initial concepts) have been adapted for use here. In particular the requirement to register multi-modal image data into a common reference frame to sub-cellular precision, has led to the development of an image registration technique based on astrometric methods commonly used in astronomy. For instance, large sky surveys require accurate registration and stitching, where the astrometric calibration relies on matching to a well known defined set of reference marker stars (e.g. stitching thousands of sky images to form a map of the Milky Way’s inner disk^6^, or a multi-epoch, multi-wavelength atlas of the Milky Way’s Bulge^7^). The technique developed here of embedding a “star field” surrounding each tissue sample, allows for efficient and accurate registration across all image data sets, as the embedded “star field” beads are visible in each imaging modality and provide a fixed reference against which positional registration and corrections for image deformations can be made.

From the scientist’s point of view, the traditional scenario where users connect to an archive or repository and download data to their own computers to perform analysis are bound to be unfeasible. We favour instead a model where users connect to a cloud based remote system close to where the data are and having available tools to further carry out analysis using the full power of the ’IMAXT cloud’. This model changes the way scientists interact with data, how they solve problems and share results with colleagues. Similar endeavours are also starting across many other scientific fields, in particular earth sciences, with the Pangeo project^8,9^ and the Planetary Computer (https://planetarycomputer.microsoft.com) and the LSST project in astronomy^10^.

The move to cloud based analysis comes however with its own challenges. Analysis pipelines are required that perform tasks with high parallelism, as well as new tools and data formats that allow chunked parallel reads and writes and are cloud storage efficient. In the next sections we describe our approach to these and other challenges and introduce some of the analysis pipelines and methods used.

This technical report focuses on the data analysis infrastructure and methods used to produce datasets that are science ready. The pipelines described here generate 3-dimensional, molecularly annotated models of breast cancer. The first three of these models are presented and discussed in our companion paper (Bressan, D. *et al*. 2021, Nature Cancer, submitted), which describes how novel Virtual Reality technologies support collaborative, yet distributed, fully immersive exploration of the data, in addition to a full description of those biological samples and the data availability.

## Results

Efficient processing and analysis of samples acquired by imaging techniques becomes a challenge due to increasingly large data volumes for each modality, increased rate at which these are produced and the variety of modalities that can be used to obtain a complete picture.

In this report we introduce the infrastructure that we have developed to address these challenges. Data analysis pipelines in the cloud allow for those pipelines to utilize resources as needed and scale when required as they run close to where the data are located. In the same vein scientists can access the full power of the cloud to perform their analysis and share results with colleagues without having to transfer large amounts of data and with dedicated software tools readily available.

Using this infrastructure we build high throughput automatic pipelines for stitching, registration and segmentation of the datasets generated by the multiple modalities that are at a later stage federated to build a complex 3D model of a tumour.

Creating a 3D STPT cube from the data obtained by the microscope is a two step process. In the first instance we stitch all the fields of view (tiles) that make each slice. In order to do so, we correct the tiles from instrumental effects and compute the offsets between all adjacent tiles using their overlap areas. Using these offsets as free parameters we find the position of each tile in the stage by minimizing the overlap residuals. Once a slice is stitched, we run it through a neural network model trained to segment the beads. The detected beads are then fitted using a high order Gaussian function to determine the centre and diameter. Since the bead diameter is larger than the thickness of the slice, there will be quite a few beads in common between consecutive slices (they will have different diameters but their centres will be accurate). Using these common beads we register consecutive slices pairwise assuming a rigid transformation.

Figure 2 shows a final aligned 3D STPT cube for 100 × 15 µm sections. For this visualisation the images are resampled to 5 µm pixels with a standard slice thickness of 15 µm. This sample was processed by an orthotopic injection of the fluorescent 4t1-E subclone^11^ shown in channel 3 (green) which is known to undergo vascular mimicry. In addition this sample was perfused with the DiI lipophilic dye seen in channel 2 (red) to highlight vessel structures.

**Figure 2:**
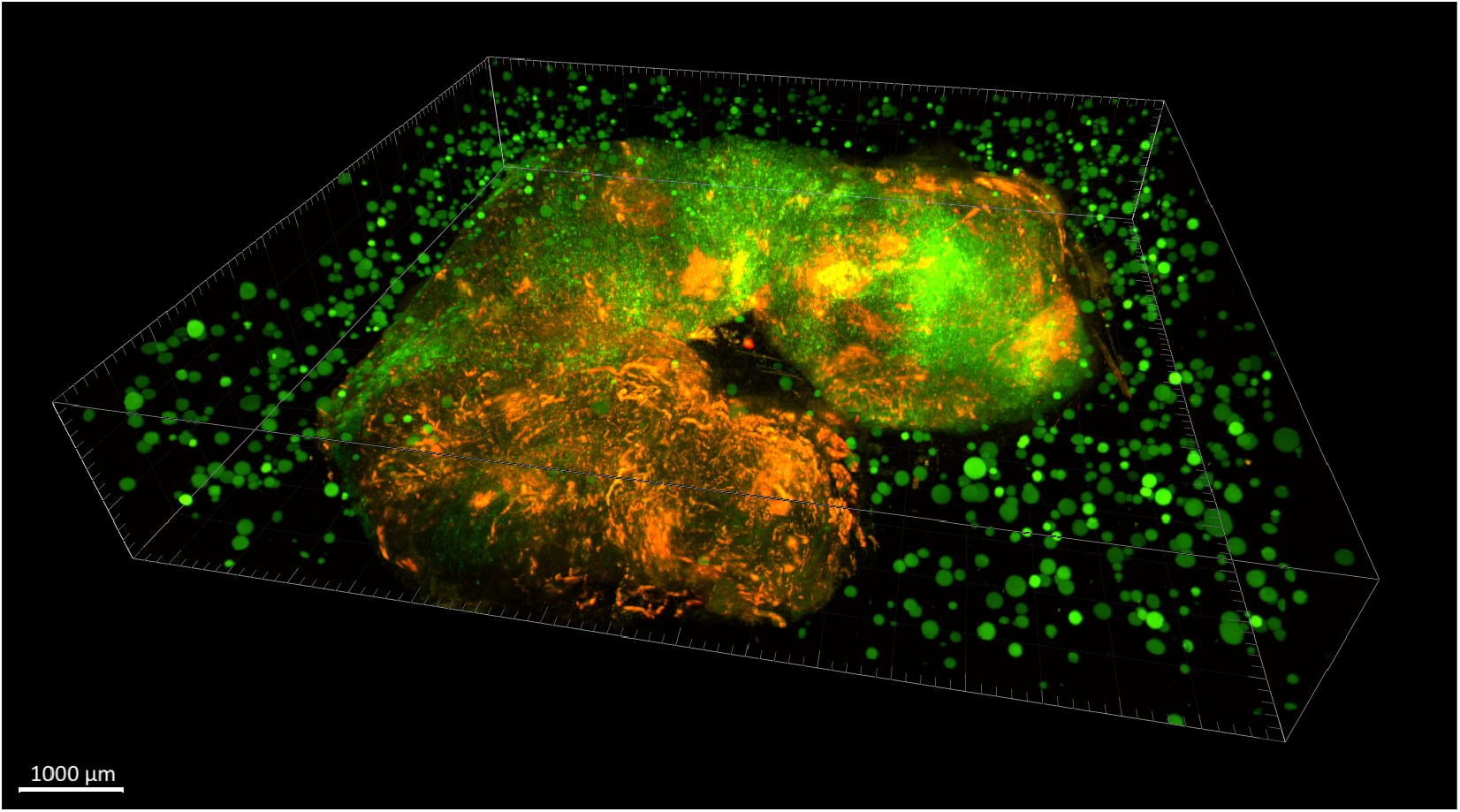
3D STPT aligned cube. 3D visualization of a stitched cube made of 100 slices, each of which made from 8 × 9 tiles. The green fluorescence beads are clearly visible in the medium outside the biological tissue and prove to be crucial for all stages of registration.

The total size of the final data cube at the original resolution is around 2 TB. The time that it takes to run the full stitching and registration pipeline and produce the final cube varies depending on the number of cores allocated for the process and the speed of the disks. The choice of resources is given by the requirement to process a sample in about the same time it takes to acquire it. Typically using 100 cores and 3 GB of RAM per core we get a processing time of 8 hours for a sample of 100 slices. Disk I/O performance is important since we compute a few intermediate files that we save to disk. Using solid state disk storage reduces the overall time by a factor of two.

The multi-modality registration software is fully compatible with STPT, IMC and low-magnification fluorescence imaging data (i.e. whole slide scanning). The software has been tested over a sample for which STPT and IMC data haven been collected over 20 physical slices, using an Axioscan fluorescence scanning microscope as an intermediate step to simplify slide identification.

There was initially a lower alignment performance between IMC and STPT, likely due to mechanical deformations introduced into the sample when collecting slices from the STPT microtome and depositing them into a slide, and also strongly influenced by the low number (*∼*10) of fiducial marks present in the field of view of the typical IMC acquisition for the test 3D datasets.

We implemented an improved embedding chemistry based on more rigid hydrogels developed by Tissuevision Inc., reaching bead densities up to 10 times higher, and have increased the overlap between the individual image tiles forming the STPT data cube, improving the bead identification. This allowed us to improve the realignment precision on individual (i.e. not belonging to a 3D sample) STPT sections and their matching IMC datasets. Resolutions allowing single-cell realignment are now attainable.

Computing times for registration between datasets are 2 minutes per slice per core, and the typical fiducial matching error is of 5 µm between STPT and Axioscan and of 8 µm between IMC and Axioscan. Figure 3 shows the result of the full registration and reprojection.

**Figure 3:**
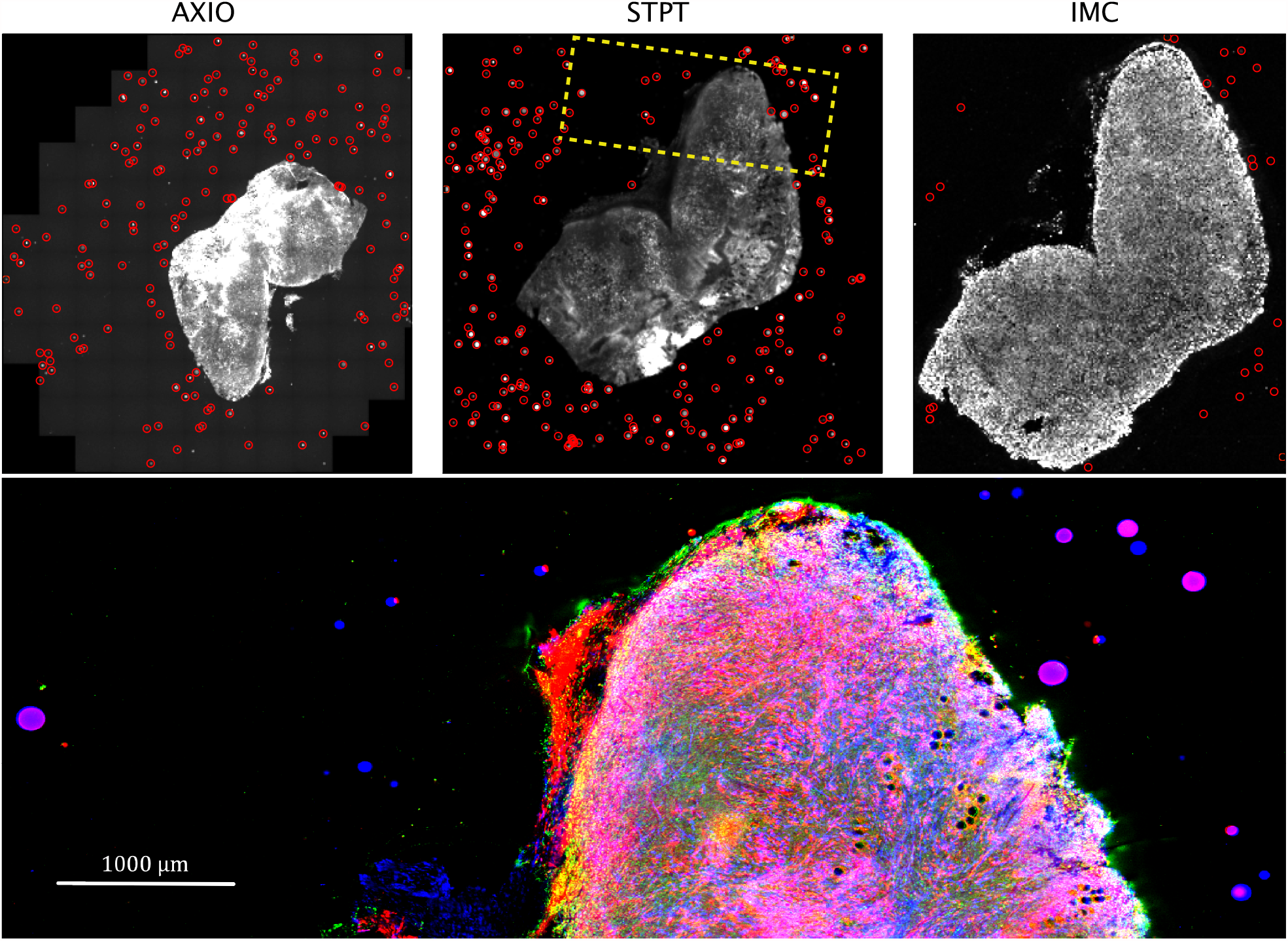
Multi-modality image registration. Registration across different modalities is achieved using the spherical fiducial beads. These are automatically detected on single-channel images, and a measurement of their geometric centre is obtained. The top row of images show the location of detected beads in an Axioscan, STPT and IMC slide. A first coarse alignment is carried out using 32x downsampled images, and with this and the fiducial centre coordinates, an affine transformation matrix is calculated. With this, we can reproject between modalities. The result can be seen in the bottom panel, an inset of the box outlined in the STPT image once registration has finished; here Axioscan occupies the red channel, IMC the green and STPT the blue.

**Figure 4:**
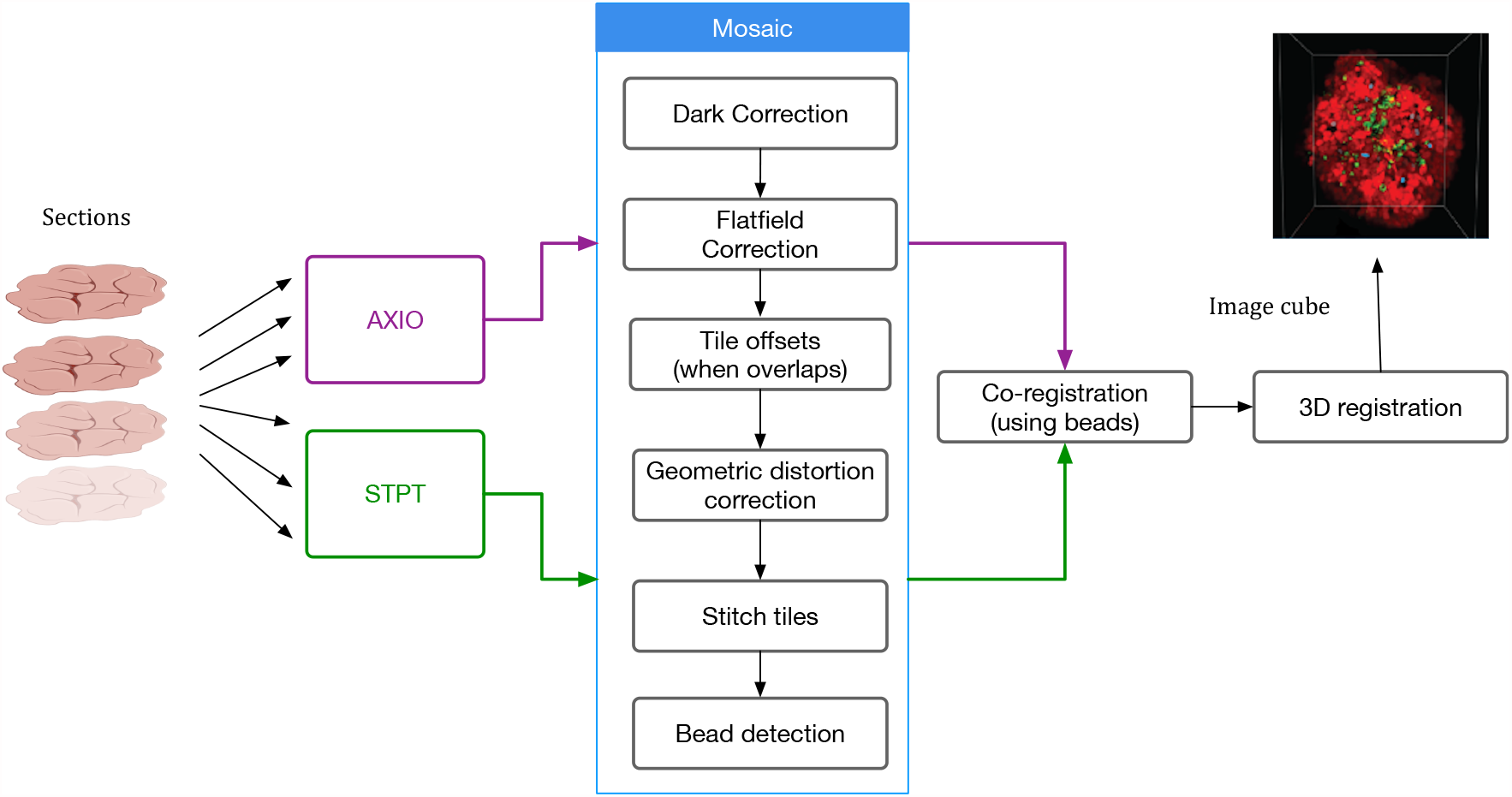
Processing pipeline steps applied to both STPT and Axioscan raw image data. Note that in the experimental flow the STPT acquires the images of the slides and does the sectioning after which they are acquired with Axioscan.

The nuclear segmentation pipeline achieves a good performance on IMC images using thresholding based segmentation. Using the iridium DNA intercalator as a marker of the nuclear channel, the pipeline extracts cell locations and relevant areas in this channel and then measures the integrated intensity over each cell’s segmented area in every channel. Our results show a high correlation of detections when compared to manual count. Some 89% of cells are correctly detected while we estimate a false positive detection rate of *∼*10%. The results have also been validated scientifically elsewhere^12^.

Our baseline May 2021 IMAXT infrastructure and analysis system release includes the following instrument specific pipelines:

- **STPT mosaic pipeline**. Performs stitching of individual tiles for each slice, taking into account overlaps between tiles and geometric distortions followed by 3D registration of all slides in a sample.
- **STPT tissue segmentation pipeline**. A tailored pipeline optimised for segmentation and reconstruction of stroma and vascular structures in STPT datasets.
- **Axioscan mosaic pipeline**. Similar to the STPT pipeline but optimised for the fluorescence scanning datasets.
- **IMC nuclear segmentation**. Performs nuclear segmentation, currently using a watershed segmentation based algorithm, and produces a catalogue of cell positions, shapes, intensities and image derived properties.
- **MERFISH mosaic and decoding**. This pipeline performs stitching, segmentation and decoding of genes^4^.

We provide a detailed description of the algorithms involved in stitching, registration, nuclear segmentation, volume segmentation and data federation in the Methods section, including the final co-registration of the IMC segmented images into the STPT ground truth reference frame where registration errors of *∼*7 µm are achieved (median error across the sample).

## Discussion

New imaging technologies and techniques are opening the possibility to explore fully molecularly annotated tissue samples at the subcellular level. The full discovery potential is realised by successfully federating the multi-pathway input data streams, such that the accurately registered three dimensional model of the tissue sample can be realised.

IMAXT is taking a holistic approach, in order to meet one of the current Cancer Grand Challenges, generating detailed maps at the single cell level of all cells and cell types in a tumour, creating 3D renderings of tumour models in which both the tumour cells and cells of the tumour microenvironment are annotated using a fluorescent code. The 3D renderings are created from samples placed in a STPT microscope. Alternating optical and physical sections, producing overlapping 3D plates are stitched back together to create the full rendering of the original sample. The physical sections then are available for further analysis; here we introduce the use of Imaging Mass Cytometry to provide proteomic annotation of the tissue cells.

The challenge that has been overcome here is in creating a robust and efficient analysis pipeline to take the digitised information from each imaging modality, and successfully integrate and align the various modalities. The registration is derived from techniques developed in astronomy, where a ground truth reference frame is defined by the marker beads embedded in the sample block. The visibility of the beads in all image modalities enables their use in several areas. The bead signature for each image section allows for the sorting of physical slices *in silico* and removes the requirement for complicated physical sorting in the STPT slice collection bath. Generating the geometry of each image from the beads and overlapping beads between individual data segments allows for effective stitching of large image mosaics. The STPT images provide a ground truth image reference frame into which all other imaging modalities can be re-projected.

The IMAXT processing infrastructure is able to integrate data from a range of image sources. The IMAXT Data Model is constructed to allow a full representation of the data to be captured in the associated metadata. Tracking of all processing steps is ensured through processing history updates to this metadata. The complete data processing software analysis infrastructure has been deployed on the IMAXT cloud, centralised on underlying hardware at the IMAXT Data Processing Centre in Cambridge. This cloud based model enables the IMAXT collaborators to interact with the processed data products through a sophisticated science platform. The basic analysis pipelines generating the instrumentally calibrated science data products are essentially fully automated and able to process the large data sets generated by the IMAXT instrumentation suite. The federated data catalogues are provided as flat files, and also through a relational database. Subsets of the full data set are seamlessly streamed to the IMAXT Virtual Reality suite of tools for rich immersive visualisation. This is described in the associated paper (Bressan, D. *et al*. 2021, Nature Cancer, submitted).

Future developments of the IMAXT analysis system will focus on the integration of additional modality specific processing pipelines to allow the integration of additional transcriptomic and proteomic measurements, for instance from MERFISH and HiFi. Currently the dissociated single-cell analysis informs the definition of the gene and protein panels used in the spatial imaging modalities. In the near future the integrated 3D annotated tumour models will also allow feedback to single cell studies, for instance relating spatial clonal evolution in time sampled Patientderived xenografts (PDX) models with tracking of allele- and haplotype-specific copy number aberrations at single-cell resolution.

This will lead to richer annotated tissue samples at the sub-cell level and provide the underpinning data for spatial’omics’ studies where the understanding of not only the biological make up at the cell level but also the spatial context is required. The IMAXT analysis infrastructure represents a significant step forward in underpinning experimental imaging advances, and coupled with IMAXT’s novel VR enhanced visualisation and data immersion tools, will lead to paradigm shifts in our understanding of tumours and their environments.

In summary we present the IMAXT data architecture including detailed descriptions of the analysis chains developed to enable the construction of accurate (to subcellular precisions), molecularly annotated, breast cancer models constructed from high spatial resolution STPT and IMC imaging.

## Methods

### Data analysis infrastructure

The requirements of a data analysis infrastructure able to process and analyse the vast amounts and variety of data for this project are: a) being able to run noninteractive data pipelines specific to each modality in an automatic fashion, i.e. as soon as data become available; b) running user analysis batch jobs with specific resources allowing for resource scaling; c) being able to perform interactive analysis that runs close to where the data are and d) running highly parallel optimized workflows. Together with these we also need in place on demand user storage, as well as data access processes and policies. The architectural approach taken has heritage in systems developed to handle optical and near-infrared imaging and spectroscopic data in astronomy, e.g. VISTA^13^ and WEAVE^14^.

#### IMAXT Cloud

The aim of the IMAXT Cloud is to allow users to run data analysis pipelines remotely and to work and analyse data interactively close to where the data are located, using already available software packages and utilising computer resources as required. The core of our analysis platform is a Kubernetes (https://kubernetes.io) cluster that runs on premises. This cloud architecture allows for the flexibility required to allocate resources and scale jobs as needed. It allows us to treat all physical computers of the cluster as one unit as well as trivially scale up new hardware, deploy isolated applications, provide dynamic resource provisioning and crucially maintain as good degree of stability and availability.

The IMAXT Cloud is deployed in our own dedicated hardware. However it is worth noting that we use similar approaches to commercial Clouds. The system could be deployed with a few configuration changes in the Google Computing Platform, Amazon WebServices, Microsoft Azure or any cloud that provides access to a Kubernetes cluster.

#### Interactive analysis

The main language used for data pipelines, analysis and infrastructure (archive, notebooks, background scripts, web services) across the project is Python^15^. Python has been acquiring great popularity across all modalities of industry and is indeed one of the main programming languages used in data science. Additionally R^16^ and RStudio^17^ are offered for interactive analysis and user batch jobs. Interactive analysis is powered by Jupyter Notebooks^18^ that are spawned in our cloud on demand using JupyterHub (https://jupyter.org/hub). User environments have a large variety of packages already installed and readily available so the user experience is as easy as to navigate to a website and be taken to a live Jupyter notebook session from where they can run interactive analyses and visualization.

A crucial point is that software environments are the same for all users, which improves shareability and reproducibility of analysis.

#### Remote desktop environments

In order to facilitate different types of data access and analysis, we also provide access to on demand remote desktop environments. This allows the user to use the cluster as a remote machine with a user specified request of resources and access to preinstalled analysis and visualization tools (Ilastik, Cellprofiler, QuPath, Fiji …).

#### Batch jobs

Batch jobs are powered by a custom job scheduler that powers automatic data pipelines and allows users to submit their own analysis to the cluster. Data analysis jobs can be submitted using a command line from anywhere (i.e. users do not need to connect to a login node) and require no experience or technical skills. More detail on batch jobs can be found in the supplementary information.

#### Parallel and distributed computing

We use Dask^19^ for parallel and distributed jobs. This allows for worker nodes that are provisioned on demand and can scale up or down depending on the workload needs. Dask also allows us to distribute large datasets across many nodes and perform efficient computations in parallel. Coupled with efficient data formats that can read and write chunks of data in parallel and independently allows for highly efficient analysis tasks to be performed on larger than memory datasets. These analysis tasks can automatically scale from a single computer to a 100 node cluster.

### Data model and data formats

The IMAXT project involves a large number of different data formats used across different modalities, some of which are created *ad hoc* for some instruments. Among the most common data formats, we find TIFF (Tagged Image File Format), HDF5 (Hierarchical Data Format) and CZI (Carl Zeiss Imaging format). This file format fragmentation is a real issue, as demonstrated by the fact that the Bio-Formats^20^ software tools incorporate plugins to read more than 150 proprietary file formats.

As for any large scale multi-modality project, it is necessary to standardise the data model and format(s) to control and streamline data handling, bookkeeping, and to make it accessible to all users.

The other component of the data model is the metadata. We define metadata as any information relating to the biological image/data that is potentially required for its scientific analysis. That is not only image related data like the dimensions and pixel scale of the image, but also the information on the instrument/microscope, the biological sample and preparation/processing of the sample. Storing all meta-data together with the data is necessary for proper bookkeeping and preventing any related human error. The metadata from different IMAXT modalities are included in separate files and/or not stored in a standard format. More importantly the information on the sample or related pre-processing is often missing and only stored by the lab users in different formats and media.

Early in the development of our infrastructure we identified that we needed a data format that allows for parallel reading and writing of different chunks of a dataset. This data format should also be able to contain both image data and metadata, to avoid any user misidentifications as data moves from one node to the other. It needed also to be flexible on metadata it can hold, as we have different modalities and instruments with a different range of metadata. These reasons lead us to choose a new emerging data format called Zarr (https://zarr.readthedocs.io). In Zarr datasets, the arrays are divided into chunks and compressed. These individual chunks can be stored as files on a filesystem, as objects in a cloud storage bucket or even in a database, making it efficient for clusters of CPUs to access the data in parallel. The metadata are stored in lightweight .json files and allows all the metadata to be in a single location which requires just one read.

All input data are then converted to Zarr, and all the metadata available stored within the dataset. All our pipelines work exclusively on Zarr datasets. We also welcome more recent developments where Zarr is the base specification for storing bioimaging data in the cloud^21^. We do as well provide custom converters from Zarr to pyramidal OME-TIFF^22^ since this is a widely supported format of many external tools.

Information about the available data, data products and their metadata is ingested into a relational database that allows interrogation via the IMAXT Web Portal (https://imaxt.ast.cam.ac.uk). Any future user updates to the metadata is tracked using versioning.

### Automatic data analysis pipelines

As discussed above, many of the imaging data requires an extra level of processing to make them scientifically usable and to extract relevant information from them, from stitching to registration to segmentation. One of our main aims has been to build data pipelines that run in a fully automatic way, i.e., once a dataset arrives to our storage it is processed without human intervention and without delay.

The IMAXT data flow diagram is shown in the supplementary information. Data taken with different microscopes are manually transferred to a specific storage location monitored by an uploader application. The uploader automatically transfers the data to the IMAXT cloud storage using Amazon simple storage protocol where they are converted to Zarr format. The metadata is also written into the IMAXT database with a versioning system that makes it possible for future updates. Once the conversion is done, the relevant data analysis pipeline is triggered automatically.

The results include metadata that allow their traceability, i.e., among others, versions of the software and pipelines used, data provenance, parameters used in the processing, etc. All this information is ingested into the IMAXT database.

### Stitching pipelines

STPT is a high-throughput 3D fluorescence imaging technique. It is similar to block-face imaging with the advantage of the images being relatively well aligned after acquisition, with measured slice-to-slice relative alignments below 20 µm. Note however that acquiring multiple slices introduces a drift in the microscope and the overall alignment between the first and last slice can be up to a few hundred microns. However, STPT has various advantages over standard blockface imaging. The tissue is imaged a few microns below the block surface, thereby limiting tissue deformations resulting from the cutting process^1^. Furthermore, by using a blade vibrating microtome instead of a milling machine, the sections can be used for further processing downstream using additional imaging modalities. The scanning operation is that of a regular two-photon microscope. A laser beam is used to excite one point in the sample, and the fluorescent photons are collected by the optics into a photomultiplier. The measured voltages are digitised and stored. Following the standard notation in astronomy, we will refer to one of these digital units as a count, measured in analog-to-digital units (ADU). Knowing the gain of the instrument allows to transform back from ADU to electrons, and from there given a quantum efficiency into incident photons^23^.

An image of the field of view of the microscope, hereafter a tile, is constructed by scanning with the laser across the sample and measuring the excitation intensity produced by the laser at each point, encoded by the microscope as a pixel, with a typical resolution of 0.56 µm per pixel and a size of 2080 × 2080 pixels. Different points in the focal plane are scanned by changing the angle of the incident beam. Because sample sizes are larger (typically, by a factor of a hundred) than the field of view of the microscope, we need to acquire several tiles in order to map the full staging area. Once a tile is acquired, the stage is moved in order to image consecutive tiles at the same optical depth. Tiles are acquired with an overlap of *∼*10% (this is a configurable parameter) to allow for stitching (see below). Once the whole sample has been scanned, the microtome cuts the top slice and the process is repeated at a deeper surface into the sample. For a typical sample used in IMAXT, 100 to 300 *×*15 µm thick slices are acquired in this fashion.

These sectioned slices fall into a bath and are collected manually and deposited onto a glass slide once the sectioning/imaging process is complete. Because this effectively randomises the order of acquisition, each slide is then imaged using an Axioscan and labelled. Crossmatching each of these Axioscan images with the STPT image cube ensures traceability and the ability to reconstruct 3D volumes from the data from other modalities. This Axioscan microscope uses a CMOS detector to acquire fluorescence imaging of the slice (again using a tiling pattern to map the full slice) in a range of channels at different wavelengths. The Axioscan however images the top surface of the sample as it is deposited on the glass slide. This introduces some degree of complication in the Axioscan to STPT matching, as the latter images are of a thin optical layer some microns deep into the sample, and furthermore when depositing the slice onto the glass slide this will be randomly done “bottom up” or “face up”.

Despite the differences between both modalities, from a data processing point of view the software needs are quite similar and thus we bundle them together in this section. In order to produce science-ready images, we need to stitch all the single tiles into a mosaic that encompasses the whole stage. Before doing this, though, we will try to remove the instrumental effects present in the images, namely: dark current, flat-field correction and optical distortion.

#### Dark current correction

Strictly speaking, dark current is associated with thermal noise generated in the detector itself that is added to the recorded signal. Measuring dark current on-sample is difficult, and dark frames are usually generated by taking images under similar conditions (exposure time, temperature, etc.) but with no light reaching the detector. When these calibration frames are not accessible, statistical approaches allow for disentangling dark current and signal (e.g. BaSiC^24^).

In the case of STPT, since the detector is a single-pixel photomultiplier, thermal noise is just an additive constant to the images (we have found no evidence of thermal drift within a single tile). Background illumination and stray light are a bigger source of contamination and can be seen in median-stacked frames (Fig. 5). This contamination is only relevant in the borders of each tile, and since we always use overlapping tiles, we can get rid of pixels with high background without loss of information. For the Axioscan, background and thermal signals are very low. We therefore don’t apply dark current subtraction to these modalities, although our pipeline has the capability of measuring and correcting these additive terms.

**Figure 5:**
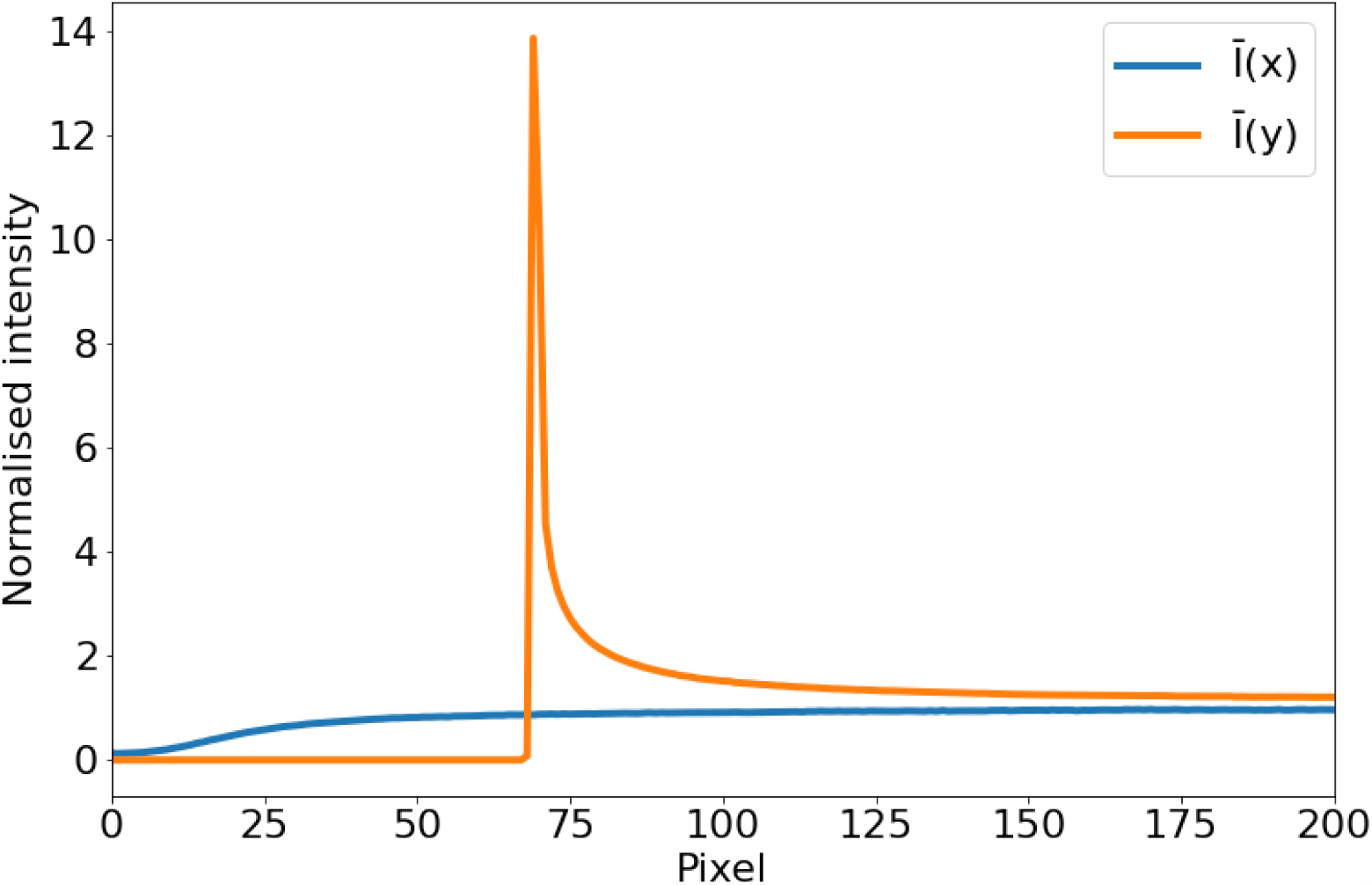
Average intensity per pixel. Normalised intensity averaged over columns (*Ī*(*x*), blue) and rows (*Ī*(*y*), orange) over the first tenth of the detector. The effect of background illumination is evident here. The first *∼* 70 pixels are already truncated by the microscope software to eliminate the most contaminated areas.

**Figure 6:**
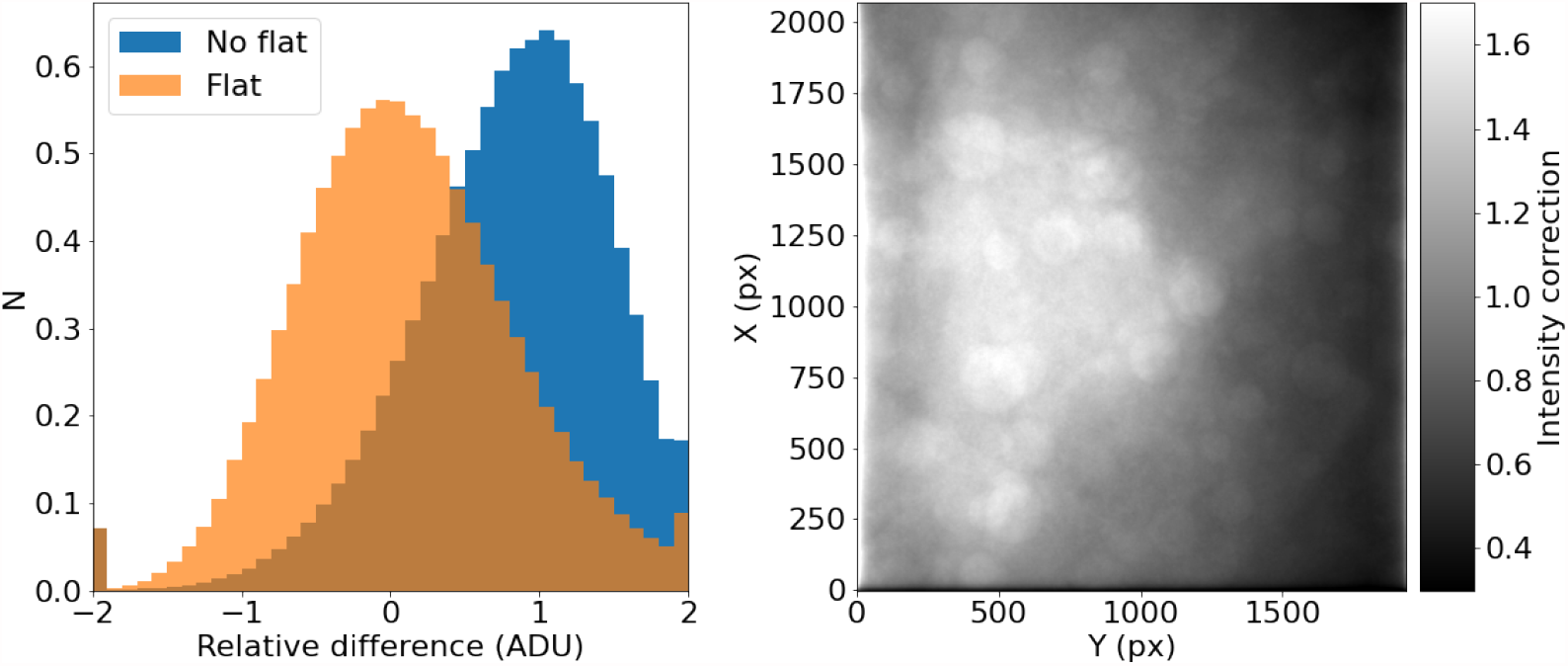
STPT flatfield correction. **Left:** Relative difference of overlapping pixels before (blue) and after (orange) flatfield correction. **Right:** Example of normalised flatfield frame.

#### Flatfield correction

In the case of most 2D detectors, the quantum efficiency is not constant across all pixels, and inhomogeneities in the lenses and other factors (like dust particles) result in a transmissivity that is a function of position in the field of view of the instrument. The standard way to measure these effects is to take the image of an uniform light source. By normalizing this image, we measure the per-pixel response function for the system at a given wavelength. Our experimental design leads to *∼*100 tiles per slice, and several tens of slices per sample. If we stack all these tiles, each pixel sees a random intensity distribution that, given enough tiles, should be homogeneous over the field of view. Therefore any variation of a location statistic like the median or the mean across the field is a measure of the different response of the system at each pixel.

Applying this correction to Axioscan tiles is straight-forward. STPT uses an unidimensional detector, so there are no pixel-to-pixel differences in response, but as the laser systematically illuminates different parts of the field of view, if effectively maps different light-paths along the optical system that can have different throughput, leading to the need to intensity correct each tile to homogenize the overall response.

It should be noted that, for a given wavelength and objective, this correction should remain relatively constant over times of days or weeks. This allows us to use this procedure with samples that may not reach enough tiles to offer reliable statistics.

#### Distortion correction

Most optical systems are subject to optical distortions. These manifest as a change of scale (resulting in a change in the shape of cells) in the field of view, being normally negligible in the centre and increasing with the distance to the centre of the field of view. This effect is particularly notorious when comparing overlapping tiles, as can be seen in Fig. 7: while towards the centre of the field of view the undistorted tiles align well, as we move towards the corners, features start to blur, to the point that they appear to duplicate close to the corners of the array.

**Figure 7:**
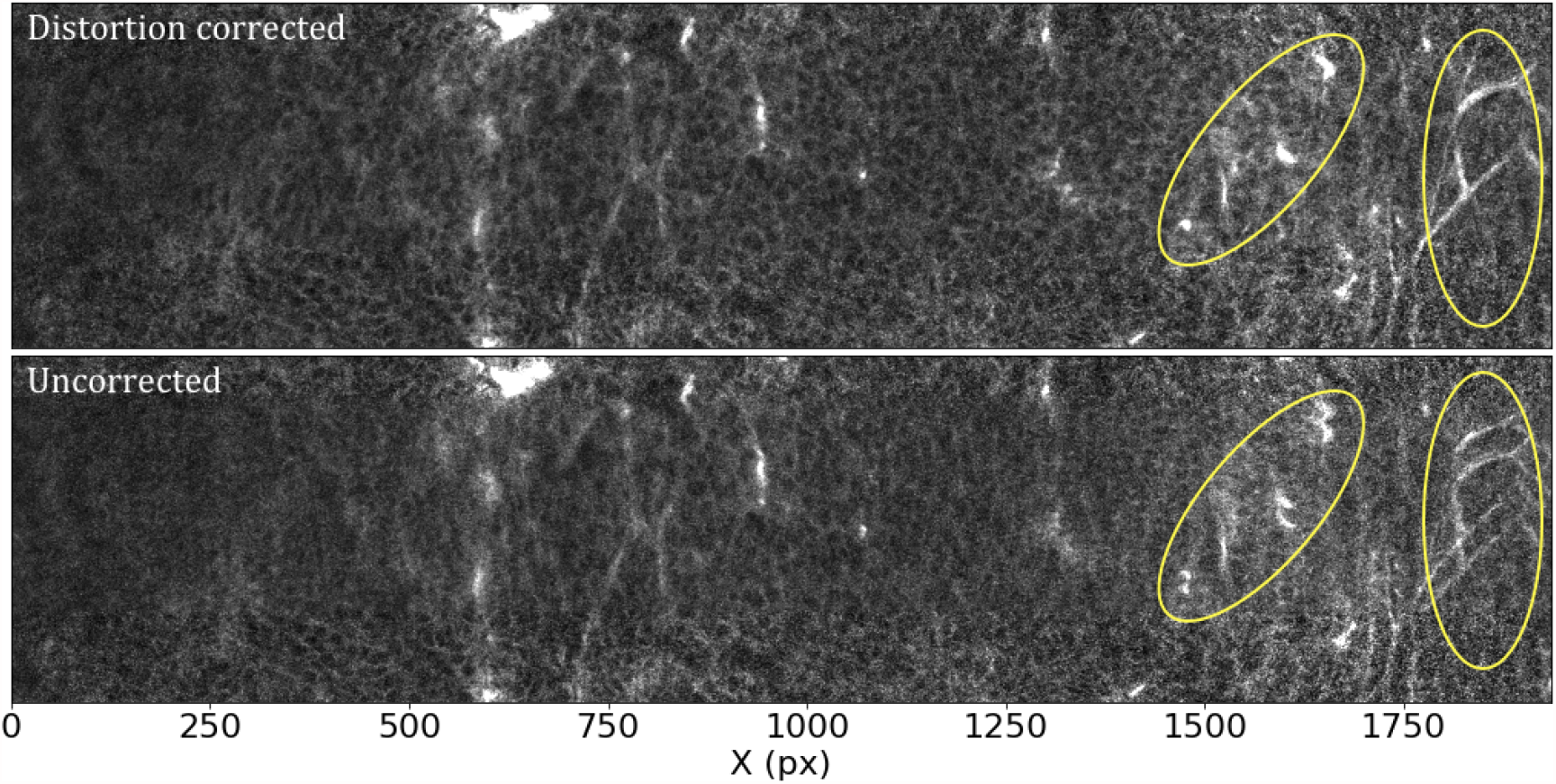
Effects of the distortion correction as applied to STPT tiles. Both panels show the same patch (500 px across in the Y direction) of two overlapping tiles. The ellipses highlight areas where distortion effects are most visible. **Top:** Distortion corrected overlap. **Bottom:** Uncorrected overlap.

In order to calculate the optical distortion of the STPT optical system, we acquired a calibration sample consisting of 10 slices in which the overlap between tiles was 50% of the field of view both in X and Y. In this configuration, each point of the sample is mapped by at least two pixels, and using feature rich parts we can minimise intensity differences between these pixels in order to fit the optical distortion of the system. The best model is a polynomial of grade 3 with different magnifications in X and Y. The resulting distortion (see Fig. 8) is a stable characteristic of the instrumental setup and used to correct the tiles for all scans of the same instrument, although it needs to be re-calibrated for each focal lens.

**Figure 8:**
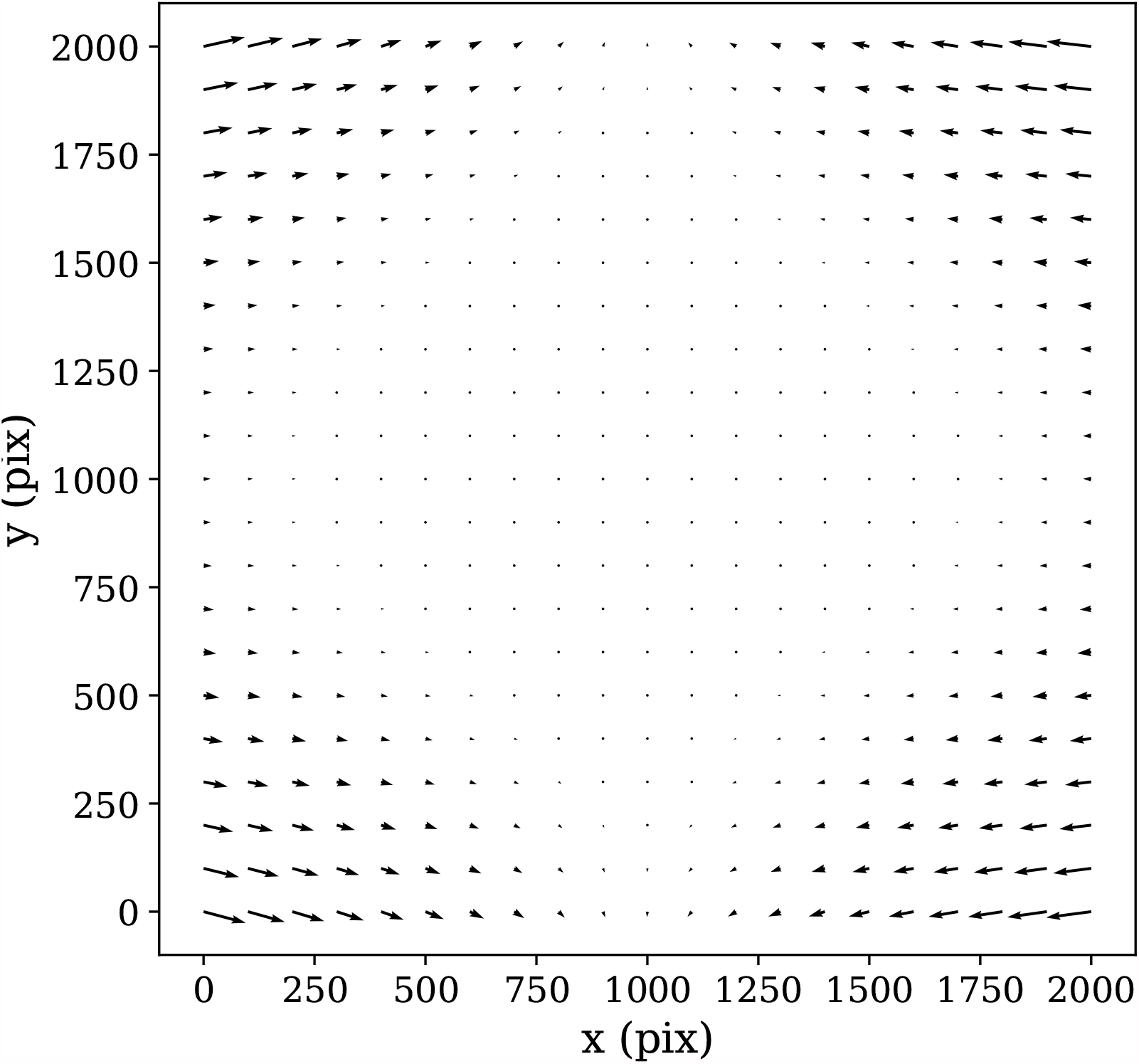
STPT geometric distortion correction. Field of view optical distortion for STPT produced by the optics of the microscope. Maximum distortion in the corners of the image is of the order of 20 µm. This is corrected before the registration between tiles by resampling the images to an undistorted pixel space.

In the case of our Axioscan, the optical distortion is small and is left uncorrected. One issue worth noting is that the microscope produces square tiles, and after processing, the pipeline uses rectangular tiles. Because applying the distortion map in Fig. 8 effectively ”squeezes” the pixels close to the corners of each tile, distortion corrected tiles have some empty pixels. In order to keep account of this, we use a construct common in astronomy called a confidence map. This is just a weight image with the same size of a tile, that encodes the provenance of each pixel. In this case, this will be a value of 0.0 for these empty pixels and a value of 1.0 for all the other ones.

#### Registration between tiles

Because the STPT microscope sees the biological samples before slicing and after minimal manipulation, imagery coming from this modality constitutes the stepping stone of much of our analysis. It is crucial that the science-grade images produced by our pipeline are the best possible and most precise representation of the sample. The control software of the microscope records the absolute position of each tile within the stage and uses this to reconstruct a raw full stage image, but we have found that there are small positioning errors that lead to a poor reconstruction of the full image. This, coupled with fact that the microscope software does not correct for flatfield or optical distortion, merits an improvement of the full stage reconstruction from individual tiles. We refer to this process as stitching.

As we have discussed before, in our configuration, STPT tiles always overlap by *∼*200 px. Starting with the recorded position provided by the microscope, we can find which tiles are neighbouring each other. Once tiles have been distorted and flatfielded, we can use overlapping pixels and find the displacement that minimises intensity differences, using as weights the confidence maps generated in the previous step. In order not to do sub-pixel resampling at this stage we also assume that the displacements are integer pixel values. This method achieves similar results as the commonly used phase correlation methods using Fourier transform^25^ but correctly taking into account masked arrays^26^. Typical differences between displacements coming from the microscope and the ones calculated on-sample are of 5-10 µm for STPT, and of around 1 µm for Axioscan. Because these differences are of the order of the expected crossmatching error, in the case of the Axioscan we use the displacementes provided by the microscope.

It should be noted that this method (as any other intensity-based crossmatch) only works where there is enough information in the overlapping regions. For pairs of tiles that have empty overlaps, we retain the displacements coming from the microscope.

Once we have the pairwise offsets between adjacent tiles, we need to compute the absolute position of the tiles in the full stage. We do this by using as reference the tile with the highest average intensity. This tile is always one that sits within the biological sample. Using the relative displacements we lay its four neighbouring tiles, and compute their absolute position. We now use each of these as reference and repeat the process iteratively until all tiles have a calculated absolute position. Because in this way there’s more than one tile-laying sequence for most tiles, we average the absolute positions weighted with the positioning error.

With these new absolute positions we calculate the stage size, in pixels, and fill this image by adding intensity values from each pixel of the individual tiles. We also generate a full stage confidence map by projecting individual confidence maps in a similar manner. Dividing the intensity image by the confidence map takes care of averaging overlapping regions, and the final science ready image is stored (Fig. 9. Note that thanks to the flatfield correction, there is no need of additional non-linear intensity blending in order to remove tile border effects.

**Figure 9:**
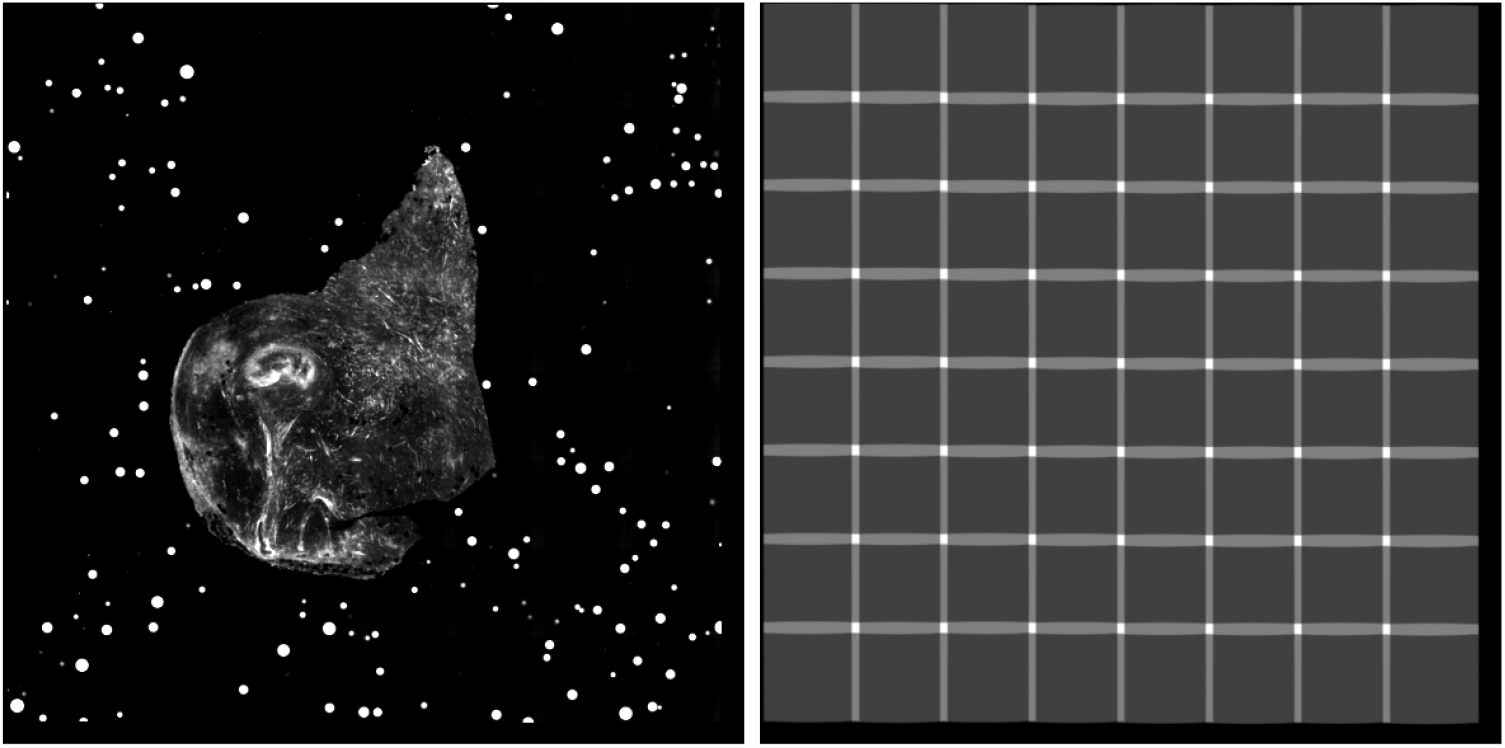
Example final stitched STPT stage mosaic (left) with associated confidence map (right). Images are padded so that all slices from a sample have the same size, hence explaining the pixels with zero confidence at the right and lower ends of the confidence map.

#### Registration between slices

In theory, the serial sections produced by STPT are inherently aligned. In reality, for each microtome pass the entire stage may shift, and some misalignment may be introduced. This can be also be due to the fact that we stitch each slice independently. Because we need to reconstruct the 3D sample as seen by the STPT microscope, we need to make sure that the relative alignment of the slices is consistent. Naively, one could think that intensity matching slices could solve this problem, but because each microtome cut removes 15 µm of sample and the STPT focuses a few microns below the sample surface, this is not possible, as images are far apart in sample depth. In order to solve this (and allow for registration across modalities further down the data processing), we introduce spherical beads in the sample. These beads have a typical diameter of 90 µm and therefore can be clearly seen in several consecutive slices. While the outline of the beads changes between slices, their centre remains constant with depth (within the natural experimental limitations of sample manipulation, microtome blade sharpness effects, etc.) and can be used as fiducial marks. This effectively transforms the circular cutouts of the spherical beads into point sources, and opens the problem to the application of a wide library of algorithms inherited from astronomy, as locating and crossmatching the position of point sources is a problem underlying many astronomical applications.

The first step in our registration algorithm is to segment the beads. Although for some sample geometries (i.e. samples where there is only one block of biological material surrounded by a field of beads, as in Fig. 9) simple thresholding and patch labelling works, for disagregated samples a more nuanced approach is needed. We have had success employing U-Net style neural networks^27^ to solve this problem, with the advantage that once properly trained, the model works in all modalities. Once we have a bead mask, we use watershed segmentation to differentiate between individual beads, and produce a label mask (Fig. 10).

**Figure 10:**
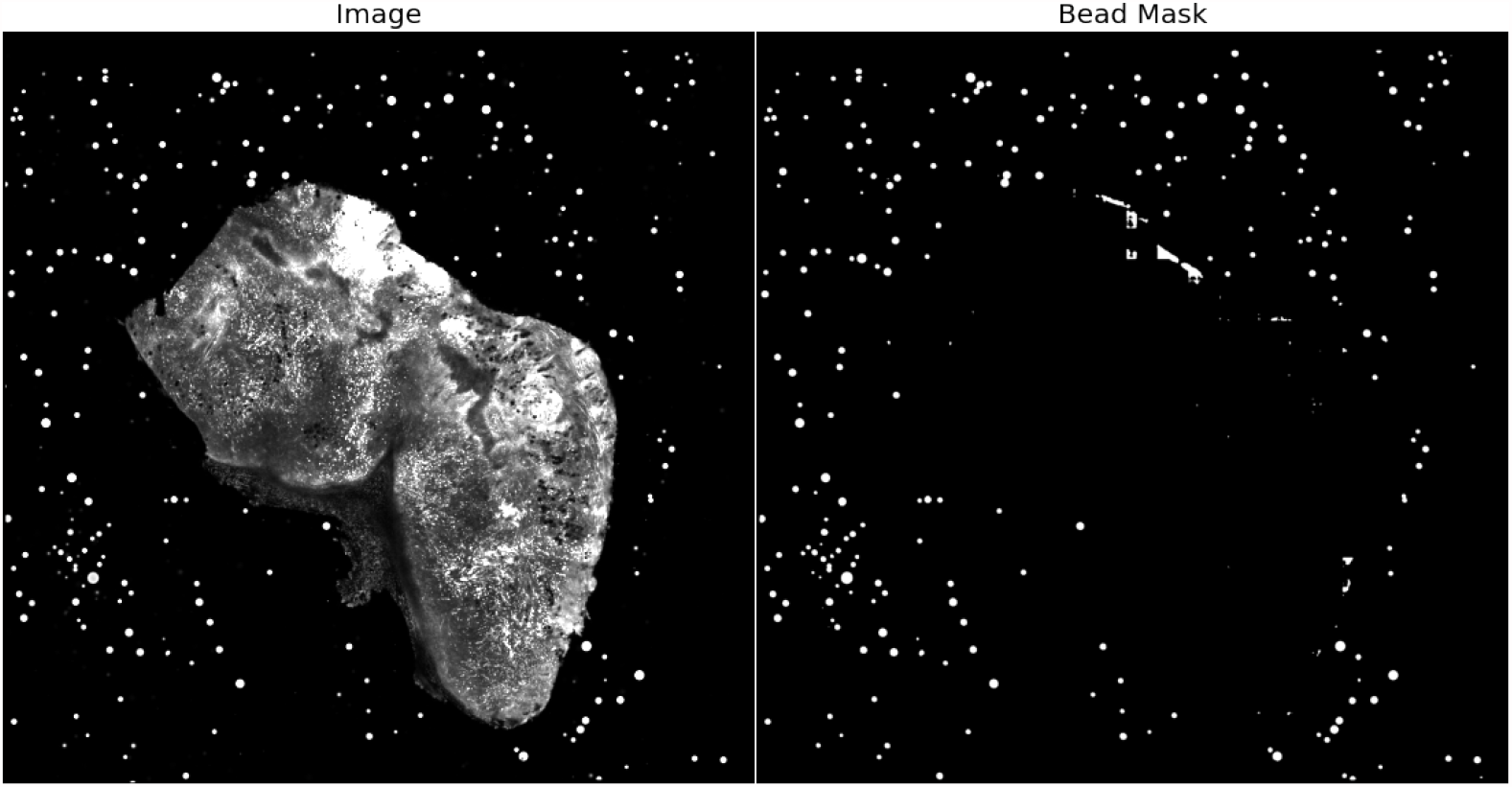
Example of an STPT mosaic and the segmented bead mask produced by a U-Net network. There are some false positives that will be removed when profiling the beads.

In order to derive the coordinates for the bead centres, we build a simple but physically realistic model of the beads consisting of:

- A sphere of constant emissivity and radius *R* centred in coordinates (*X, Y, Z*) as measured with respect to the 3D sample block.
- The STPT laser excites a layer of thickness *t* of the bead at a given depth *l* into the sample, so that the emission from this layer in local coordinates (*x, y*) is 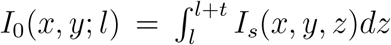. In this case, as we assume emissivity is constant over the sphere, *I*_*s*_(*x, y, z*) is just the density profile of a sphere with constant density in Cartesian coordinates.
- The depth *l* can either be in the interval [*Z − R, Z* + *R*] and so the optical surface intersects the bead, or *l ≤ Z − R* in which case we have a fully embedded bead emitting just under the optical surface. If *l ≥ Z* + *R*, the bead is between the optical surface and the observer and therefore not visible.
- The sample substrate has an optical depth *τ*, and the emission from the optical surface decays as 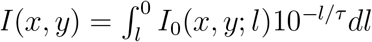.
- By integrating this last expression we obtain the 2D brightness profile as a function of (*X, Y, Z, R, l, t, τ*), that we fit to the observed profile assuming Poisson statistics.

Although this prescription may seem too complicated, it accommodates well the variety of bead brightness profiles observed in our samples (Fig. 11). For the purposes of registration, the most relevant parameters are (*X, Y, R*). The coordinates of the bead centre are the basis of our registration, and the bead radius is a simple threshold to use when crossmatching beads.

**Figure 11:**
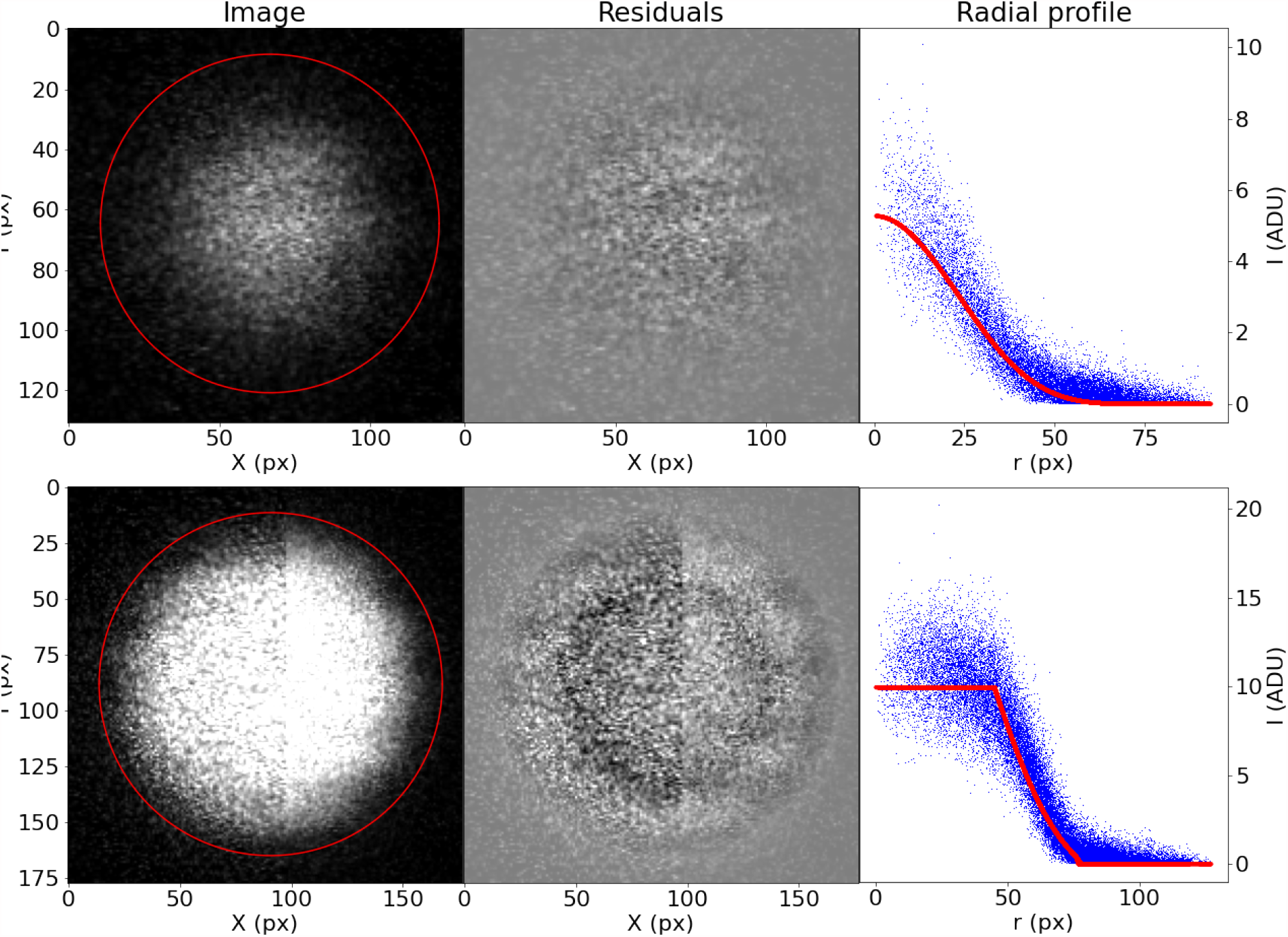
Examples of the profiling function applied to measured STPT beads.

Once we have the (*X, Y, R*) catalogue for all beads and all slices, we can proceed to the slice-to-slice registration. Because inter-slice displacements are expected to be small, for each pair of consecutive slices we crossmatch the bead catalogues by using a simple nearest-neighbour search. With all the crossmatches, we identify unique beads. Normally, due to their thickness, each bead will appear in a handful of slices. Filtering out single detections removes the vast majority of spurious contaminants.

With all the detections of a single bead *i* over all slices, we can compute the matrix with all the pairwise differences in coordinates, 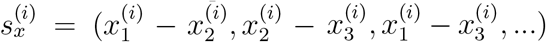, with 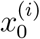 being the fitted *X* coordinate for the centre of bead *i* in the first slice, and so on. Accumulating all beads and all slices, we build the vector *S*_*x*_. We can relate this sample vector with the vector containing the absolute offsets for the slices *D*_*x*_ = (Δ*X*_1_, Δ*X*_2_, ..)^*T*^ by means of a coefficient matrix *C*:

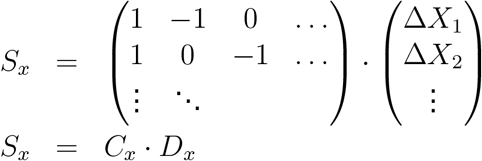

And

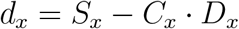

We can obtain the absolute displacements Δ*X* by minimising the equation

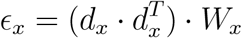

with *W*_*x*_ being a set of weights derived from the error vector for the pairwise differences in *X* coordinate for the centres. Although *C* can be a relatively large matrix, it is sparse, and therefore this minimisation is computationally efficient. The same scheme is applied to Δ*Y* (we assume displacements in both axis are independent) and from (*D*_*x*_, *D*_*y*_) we can obtain the full 3D registration of the STPT cube. An example of derived (*D*_*x*_, *D*_*y*_) can be seen in Fig. 12; while *D*_*x*_ remains more or less constant, a clear drift can be observed for *D*_*y*_. this is likely related to the effect of the microtome, that always sections the sample along the same direction, possibly causing small displacements.

**Figure 12:**
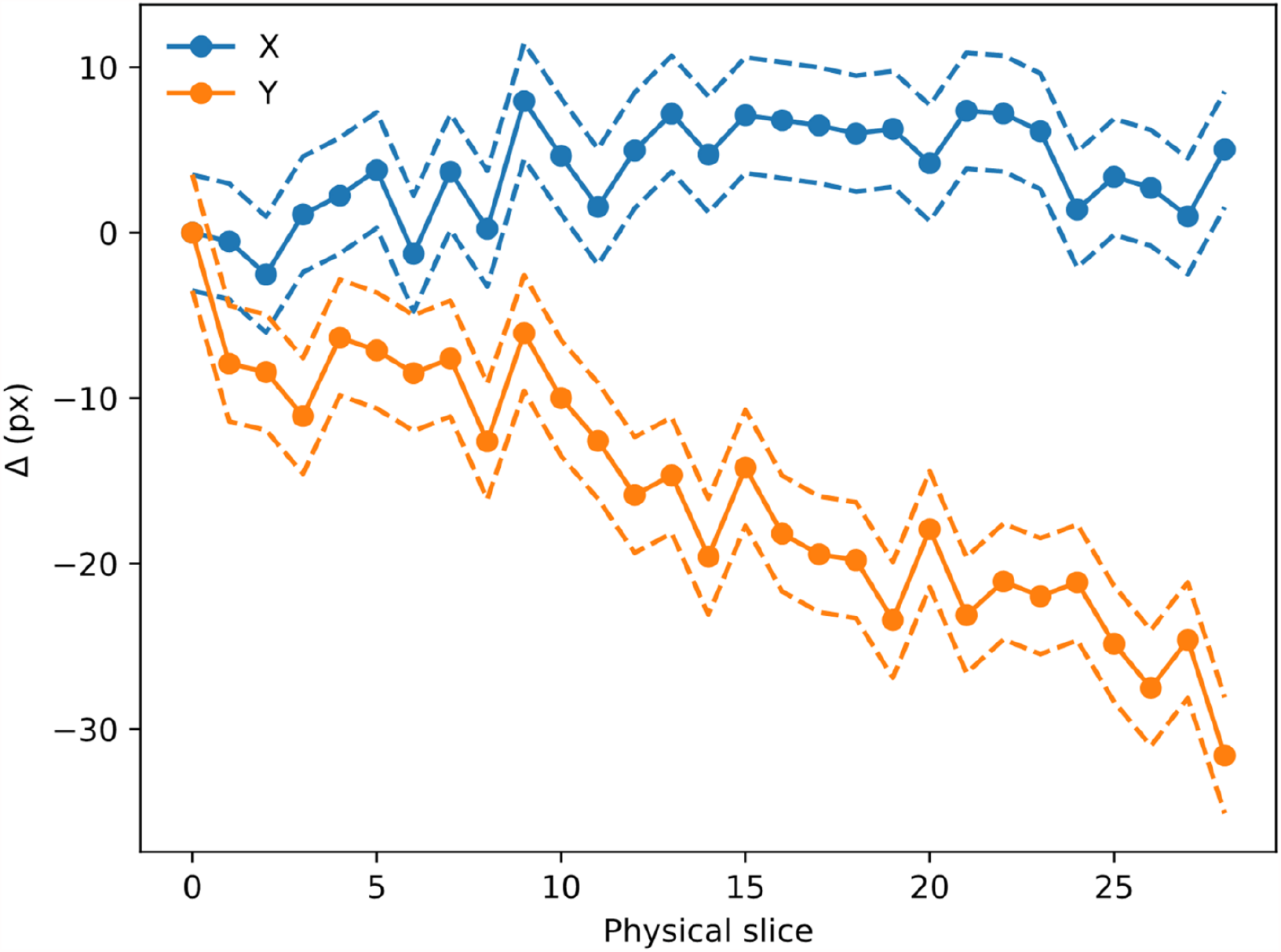
Evolution of (*D*_*X*_, *D*_*y*_) with slice number (i.e. depth along the sample). Dashed lines mark the 1*σ* error boundary. As can be seen, there is a drift in one of the directions, likely to be related to the effect of the microtome blade pushing into the sample cube. The scale for this sample is 0.56 µm per pixel.

The (*D*_*x*_, *D*_*y*_) translation values are stored in the metadata of the Zarr file and are read when processing the STPT images.

### IMC Segmentation pipeline

Imaging Mass Cytometry (IMC) is an approach to high-multiplex imaging and single-cell protein analysis^2^. The method delivers unprecedented insight into the tissue microenvironment with single-cell resolution. It helps to deeply characterise the complex and diverse tissue microenvironment by gaining an unparalleled understanding of the spatial relationships between a multitude of cell types and the role of cell phenotypes in the context of disease. It also helps to uncover novel therapeutic targets through the discovery of new biomarkers. In IMC, tissues are stained with a panel of isotope-labelled antibodies. Stained sections are laser ablated at 200 Hz at subcellular resolution, and liberated isotopes are detected with a mass cytometer to yield images quantifying the abundance and location of the proteins of interest at 1-micron resolution simultaneously. The output of this process is a data cube consisting of several layers where each layer is associated with a protein. In addition, all layers are aligned. Thus image registration as part of a pre-processing step is not necessary. The name of each protein and the order of its corresponding image layer is stored in the data cube metadata. In addition, there are few extra image layers related to the instrument calibration.

IMC images are analysed using a pipeline developed for automated, high throughput analysis of biological images. The pipeline code is written in Python and uses the OpenCV^28^ library, i.e., an open source computer vision and machine learning software library written in C++. Following reading the IMC data cube, which consists of multi-layer grey level images, the code identifies the nuclear channel from the information provided in the image meta-data (typically the channel with iridium DNA intercalator). This channel exhibits the strongest signal among all the other IMC channels and as such it serves as a reference image to identify and segment cells during the rest of the process. Though the pipeline is automated, there are a few parameters to be set before running it. These parameters are set to optimise the pipeline performance on the nuclear channel which is adopted as the reference. Therefore information retrieved for a cell from all the other channels (e.g. pixel intensities inside/outside the cell nucleus) are based on the analysis of the nuclear channel. For small tissue sizes, the IMC produces a data cube for each slice it scans. However, if the tissue size undergoing IMC scanning is large (e.g., *≥*1 mm), it may not be possible to scan the sample in a single run (for technical reason) or we may not be interested to scan the whole sample but rather individual areas of interest. In this case, the scanning area is divided into multiple region of interests (ROIs) where each ROI is processed independently and in parallel. If ROIs belong to the same tissue slice, we merge the output catalogue of the segmentation analysis at the end of the process using information stored in the metadata to convert each ROI coordinates to the stage coordinates.

#### Pre-processing

The pipeline pre-processes the reference channel before segmentation. Though the IMC data consists of 16-bit images enabling the instrument to map a high dynamic range of pixel values, the distribution of pixel values associated with the observed tissue mostly populates the low-intensity domain of the available dynamic range, i.e. not taking full advantage of the dynamic range for a 16-bit depth image. Therefore the first step in pre-processing is to enhance the reference image (e.g., the nuclear channel). To do so, we apply a linear scaling function to the image pixel values to improve their contrast by stretching the range of intensity values to span the desired values for a 16-bit image. This technique is called image normalisation and is different from histogram equalisation, where the scaling is non-linear. Next, we apply a Gaussian filter with an appropriate kernel size to remove individual high signal-to-noise pixels in the image.

#### Segmentation

To segment the image and separates the regions of the image corresponding to objects in which we are interested from the background, we apply an adaptive thresholding method to remove background pixels. The output is a binary image where pixels belong to the regions of interest and those belong to the tissue are white. It leaves us with either individual cells or those having overlaps with one another. To deblend overlapping cells, we find the coordinates of local peaks (maxima) in the binarised image. Then, we use a watershed algorithm to segment the image^29^ (see Fig. 13).

**Figure 13:**
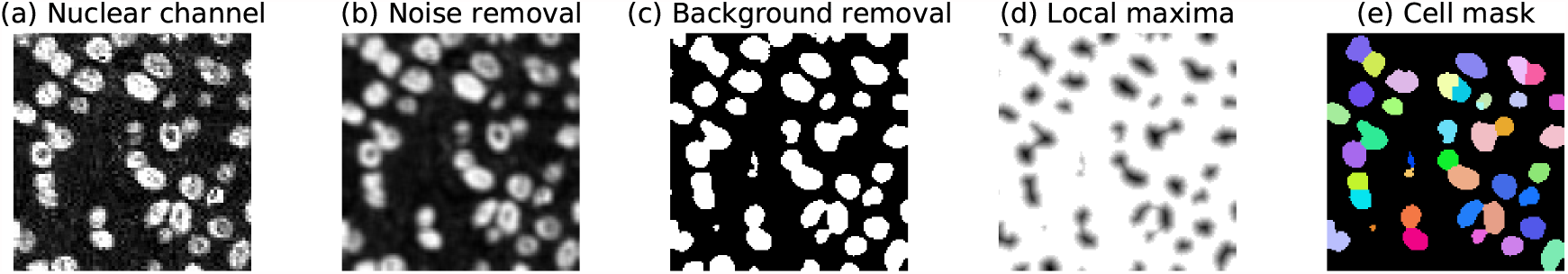
Example workflow of the IMC segmentation pipeline. (a) Initially the pipeline extracts the nuclear channel from the IMC data cube and uses it as a reference image for the subsequent segmentation process. (b) Next, we remove noises by applying a Gaussian filter to the reference image. (c) The algorithm separates background pixels from foreground pixels by applying an adaptive thresholding method to the de-noised image. The output of this process is a binary image. (d) To de-blend touching / overlapping cells, we find local maxima on the binary image and create a distance matrix. (e) Finally, we segment cells using watershed algorithm and create and image mask where individual cells have a unique id. By using the image mask, we calculate shape, geometrical, and intensity parameters for the cell nuclei and their peripheries across all available IMC channels.

#### Feature extraction and output products

Following the segmentation, we create two image masks for each detected cell. One is associated with the segmented cell itself, and the other one with the cell periphery, i.e., the two-dimensional zone surrounding the cell. For each detected cell, the code uses the first mask to compute mean pixel intensities inside the area of cell nuclei and the second mask to compute mean pixel intensities associated with cytoplasmic / membrane areas across all available IMC channels. Next, we estimate image moments for each cell and several other parameters to describe the shape and geometry for each detected cell. Finally, the code exports a table where each row represents a cell with all extracted parameters as feature columns. In addition, the code exports a mask image of all detected cells for single-cell analysis.

#### Performance

The performance of a segmentation algorithm depends on several factors. For instance, the tissue preparation method (e.g. frozen vs Formalin Fixed Paraffin Embedded), its type and thickness and the scanning resolution all affect the accuracy and performance of a segmentation algorithm. These are apart from other factors such as fine-tuning the input parameters of an algorithm or hyper-parameters of neural-net based models. Therefore the best way to assess the performance of an algorithm is to test it on specific data that is subject to the segmentation analysis. To measure the detection accuracy, we select ten random image tiles, each 100×100 µm, from the 3D STPT-IMC dataset. These tiles come from three different slices are chosen such that they represent regions of low and high-intensity pixel distribution. We count the cells inside each tile and record their positions. Then we run the IMC segmentation algorithm on these tiles and check the spatial distribution of detected cells against those selected visually. A typical cell in our sample has a diameter of 10 µm. Therefore to find the correct match between the position estimated by the algorithm and the one picked up manually, we consider a maximum matching radius of 5 µm.

The automated detection shows a high correlation (R = 0.79) with the manual count. Note that both true-positive (TP) and false-positive (FP) have contribution in the automated detection. The result shows that about 89% of cells are correctly detected (TP; Fig. 14). Though we set the matching radius to 5 µm, all TP detection are matched within 2.4 micron distance from the visually selected centroids, with a mean distance of 2.03±0.07 µm across all test samples, revealing the accuracy of the pipeline in finding the correct centres of cells. We find also that around 8-13% of detection are FP. The latter is associated with (a) the background noise or (b) multiple detection of the same cell. It is worth noticing that each image tile in our test sample, contains about 50-60 cells. Given the size of each image tile, this shows that the test sample represents a more or less dense configuration of cell distribution which increases the likelihood for false-positive detection. As such, the reported accuracy is a lower limit and expected to be higher in less populated regions.

**Figure 14:**
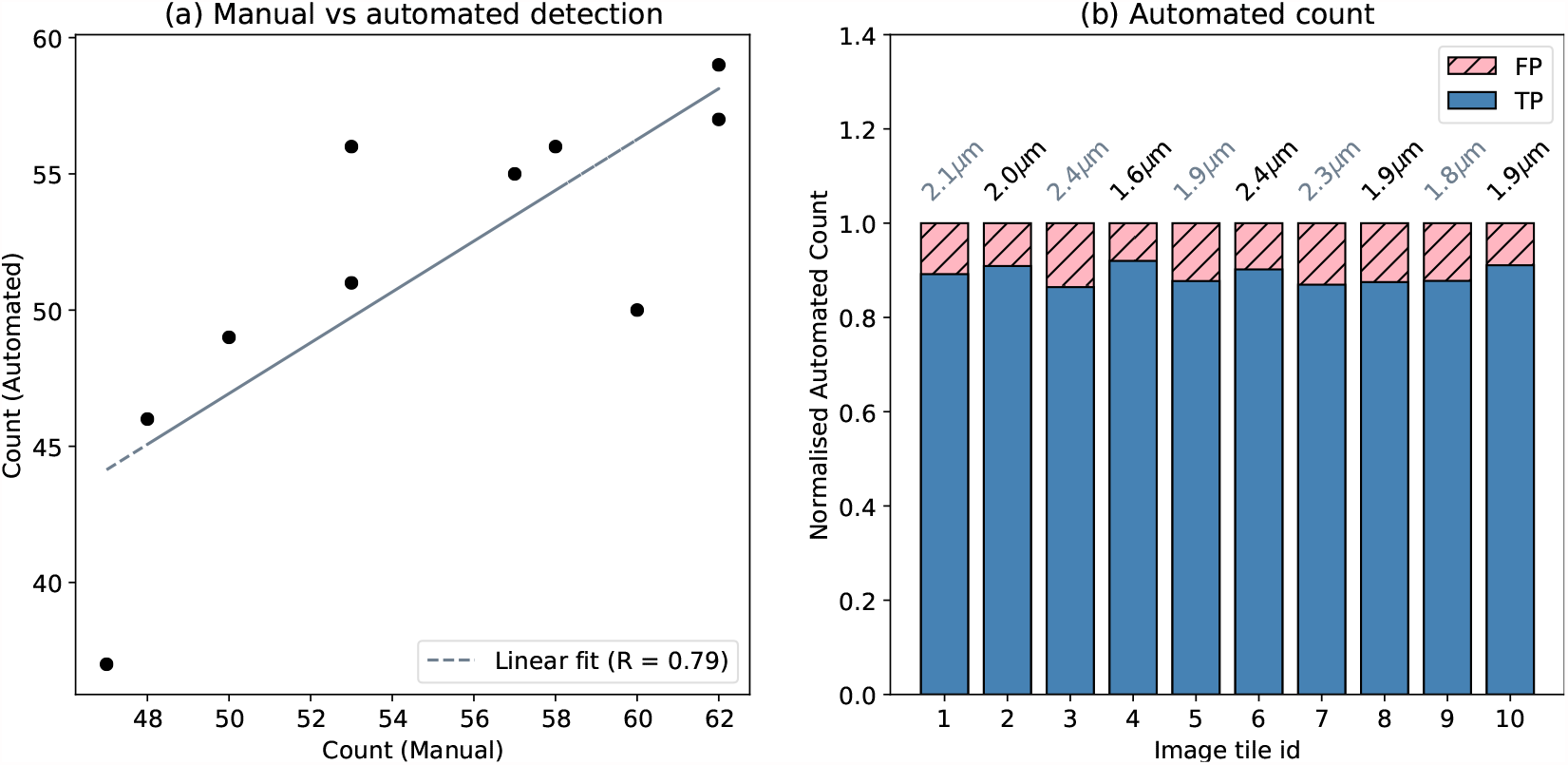
Measurement of the IMC pipeline detection accuracy, on a test sample of ten image tiles, each 100 × 100 µm, from the 3D STPT-IMC dataset. (a) Correlation of the manual count vs automated count reveals a tight correlation (R = 0.79). (b) Contribution of true-positive (TP) and false-positive (FP) to the automated detection. The y-axis is normalised to 1.0. TP detection are those detected automatically within 5 µm radius of manually selected cells. The TP contribute to 89% of the overall automated detection. The number above each bar, shows the mean matching distance in micrometres where for automated detection, matched sources have been found from visually selected sources.

As a preliminary test, we run the pipeline on a sample of breast cancer patient-derived tumour xenograft (PDTX) that was previously analysed with mass cytometry (MC). Results as analysed using the current IMC pipeline, successfully reveals the spatial distribution of cell phenotypes in xenografts as observed with MC data. The study finds that centroids of each cell cluster computed per PDX model on the MC training data shows a high correlation (*ρ* = 0.67) with the corresponding centroid following cell segmentation and classification using the IMC pipeline^12^.

### STPT tissue segmentation pipeline

While IMC is suitable for cell segmentation, the STPT modality is particularity suitable for visualising the various tissue structures, such as the stroma and vasculature. To be able to quantify these structures, several tool have been developed.

#### Section resampling

The STPT images are loaded at the level corresponding to a 4 times downsampling. They are subsequently resampled to a pixel size of half the slice spacing, i.e. 7.5 µm, while applying the x- and y-translations which were calculated during stitching and registration.

#### Upsampling

In the current setup 15 µm sections are acquired. This may cause some discontinuity between sections for oblique tissue structures, as is the case with blood vessels. To resolve this we can apply an up-sampling based on an intensity based deformable registration. With a pixel size set to half the slice spacing, a linear interpolation is performed between each pair of neighboring sections along the direction calculated by the registration. In this way we generate interpolated slices throughout the volume. This increases the out of plane resolution and improves the overall resolution when converting to a volume with an isotropic voxel spacing, which is required for some of the measurements.

#### Segmentation

Segmenting the tissue structures is done using a manually defined threshold value resulting in a binary segmentation. This is followed by a smoothing, which is performed separately for the in-plane and out-of-plane direction due to differences in the signal to noise ratio between the two. Finally, a connected component filter is applied to remove the small structures, which can be considered noise.

#### Quantification

Once the tissue is segmented, several structural parameters are extracted. These include the tissue ratios, the tissue thickness, the fractal dimension and the connectivity density.

### Data federation

While STPT process the whole sample sequentially, other modalities like Axioscan or IMC work over single slices. This implies that slice-to-slice registration may not be possible for these modalities, and in order to recover the 3D position of data from them it is required to relate them to the STPT data.

As discussed previously, physical slices are recovered from the STPT microscope in random order and deposited onto glass slides that then go through the Axioscan. The first step then in our multimodal registration is to find the best Axioscan-STPT pairs, so that we can assign a Z coordinate (i.e. depth in the sample) to each Axiocan slide image. To do this, we rely again on beads as fiducial marks. The procedure to detect/profile beads on Axioscan images is the same as for STPT.

We need to find the best correspondence between the catalogue of (*X, Y, R*) for a given Axioscan image and the (*X, Y, R*)_*i*_ for all the STPT slices. There are a few things we need to keep in mind:

- The slices can suffer mechanical deformation when being deposited onto the glass slides, leading to imperfect matches between Axioscan and STPT.
- The orientation of the slice on the glass slide is random, leading to possible left/right and top/bottom inversions.
- Not all Axioscan images may have a good STPT match. Because the first STPT image will be taken at some depth into the sample, the microtome may cut above this layer, leading to an orphan physical slice.

To address these issues, firstly we will search for a more complex transformation than the simple displacements used when registering STPT slices. We settle for an affine transformation:

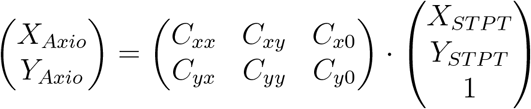

Secondly, we jumpstart the algorithm with a coarse intensity match between a 32× downsampled Axioscan image and the median STPT image for all the slices, also downsampled. This will give us the left/right and top/bottom relative orientations, and an initial estimation of the matrix of coefficients *C*_*AS*_. For each STPT slice we refine this matrix by finding the coefficients that maximise the number *N* of bead matches (*X, Y*)_*ST PT*_ → (*X, Y*)_*Axio*_ within *R*_*Axio*_. *N* should increase monotonically with STPT Z until it reaches a maximum for the most similar slice, and then onwards it should decrease as we move away from this slice. In reality (Fig. 15), *N* is a noisy function of *Z*, and so we fit a smooth function (a simple Gaussian) and find the *Z* value closest to the function maximum.

**Figure 15:**
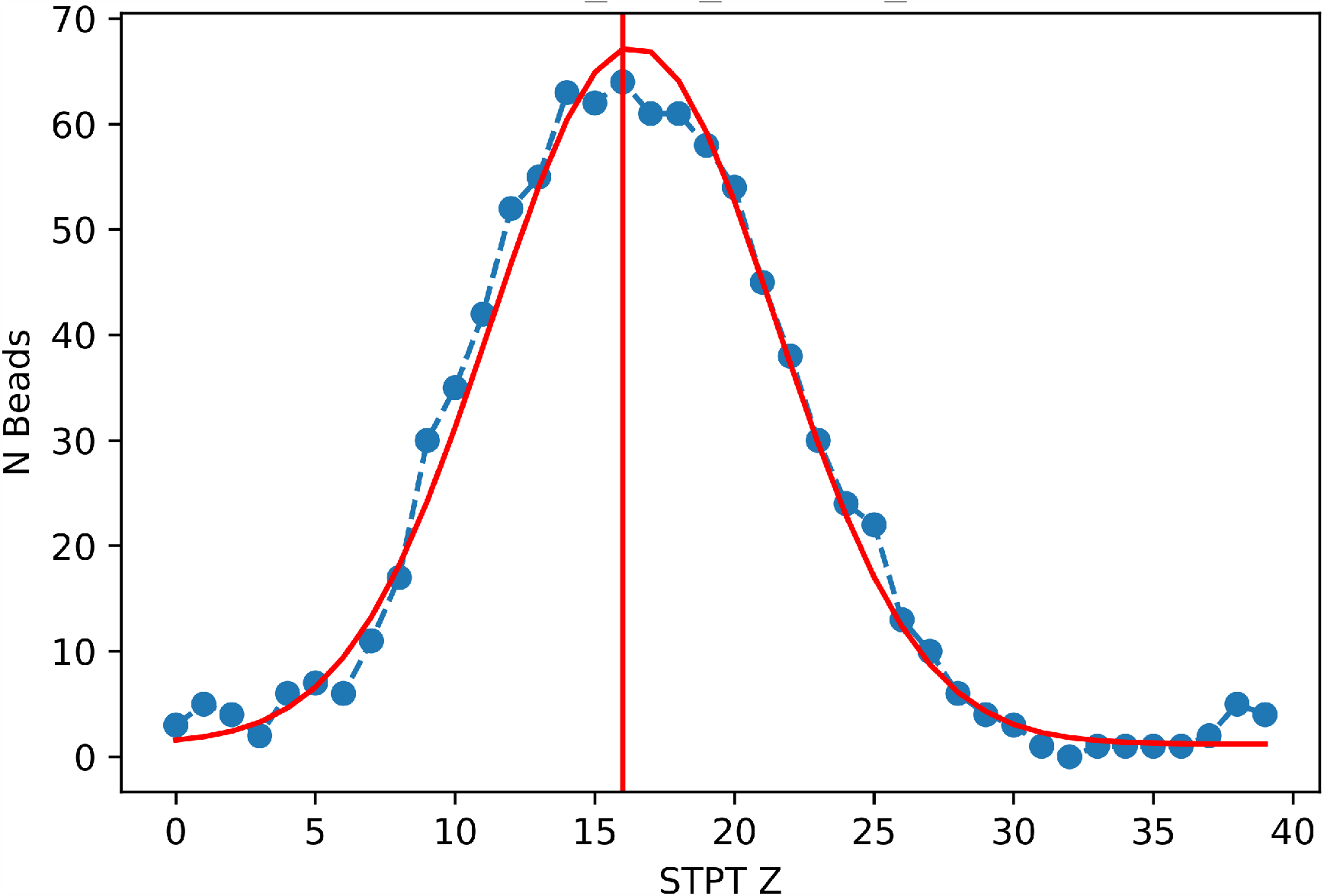
Number of common beads between an Axioscan and all the STPT slices taken from the parent sample cube. The red line is the best Gaussian fit to the evolution of *N* with *Z*, and the vertical line marks the predicted *Z* for *max*(*N*)

The result from this algorithm is the best matching STPT slice along with the corresponding affine transform between both pixel coordinates.

Once the slices are scanned they are given an identifier that simplifies further registration. Some of these slices will go to IMC, and these need to be registered back to STPT too. We use Axioscan as a sort of man-in-the-middle between IMC and STPT. Because once the slices are deposited onto the glass slides they are more or less stable^*^, IMC to Axioscan registration is quite easy, despite the fact that normally, due to the time cost involved, the portion of slice sampled with IMC tends to be small (and therefore to have few beads). Beads are detected on the IMC data in a similar manner as detailed before, and because we know which Axioscan slice correspond to which IMC, we just need to crossmatch (*X, Y*)_*Axio*_ to (*X, Y*)_*IMC*_ and obtain the coefficients *C*_*IA*_ associated with this transformation.

With this new matrix we can compound the transformation

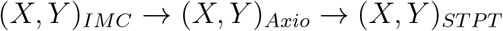

and obtain a first estimation of *C*_*IS*_; with this first crossmatch between IMC and STPT through this route, we can refine the transformation and refine the coefficients of *C*_*IS*_. We estimate the error of this procedure through the Cartesian distances between STPT beads and the respective reprojected modalities. The results are sumarised in Fig. 16; median error for the IMC to STPT registration is of 6 µm, while this figure is of 7 µm for Axioscan to STPT. These differences are mediated by the number of visible beads and the native pixel size in each modality.

**Figure 16:**
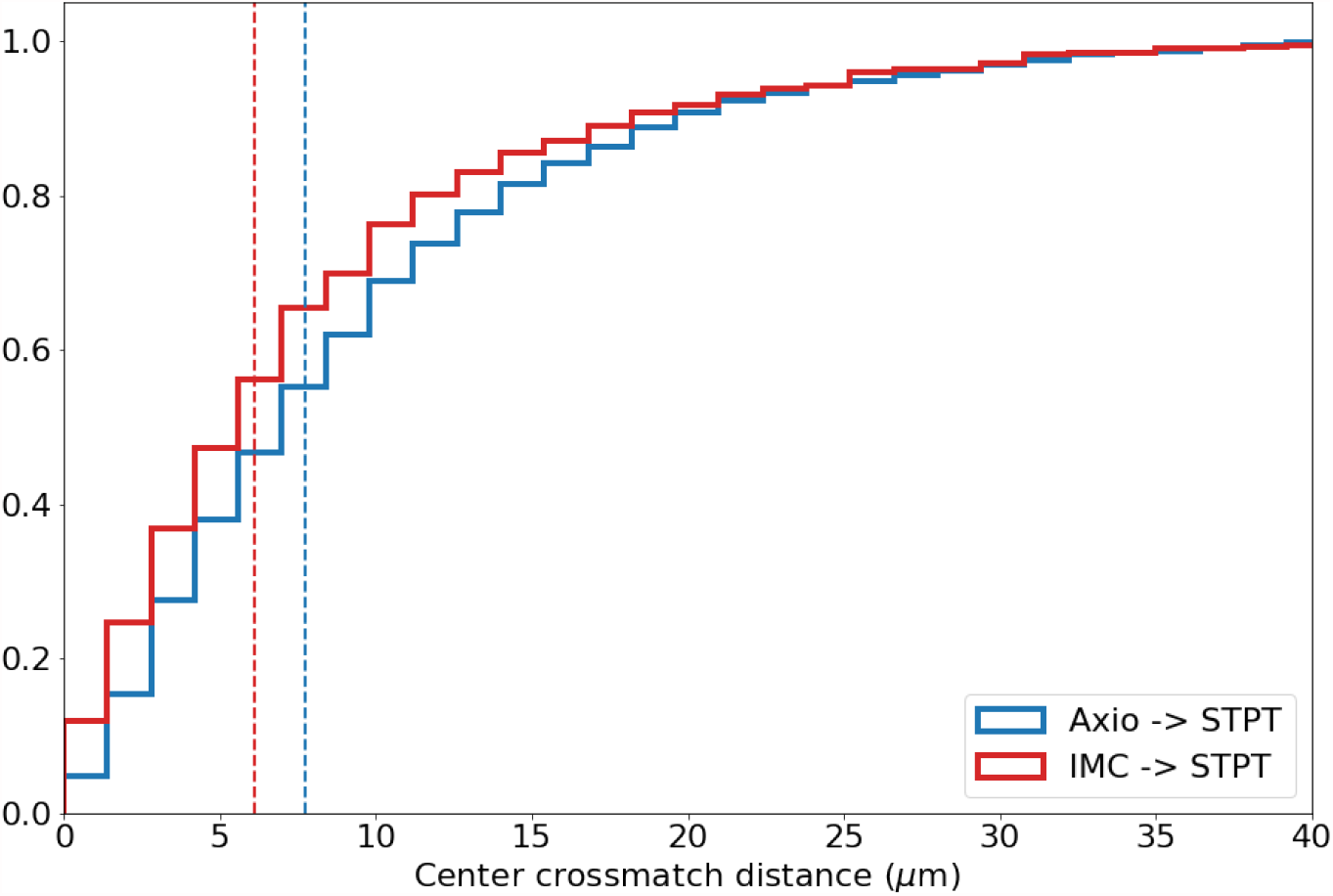
Cumulative distribution for the registration errors as measured through the reprojected centre coordinates.

A model with 6 degrees of freedom like the one we propose here may appear to be simple, but it works to sub-cellular precisions over samples with sizes of 1 cm^3^. These affine transforms have the added advantage that are easy to encode, all the coefficients have physical interpretation (rotation, scale, shear) and once we have the *C*_*AS*_, *C*_*IS*_ and *C*_*IA*_ matrixes, it is possible to reproject segmentation catalogues and masks from information-dense modalities like IMC over time efficient imagers like STPT.

An underlying assumption for this registration is that the biological sample and the beads behave in a similar way. As can be seen in Figs. 9 and 10, the beads surround the sample, but there are no fiducial marks inside the sample itself. One of the implications of this is that the errors in Fig. 16 are likely a higher envelope of the real registration errors. The STPT mosaic is built by intensity matching tile overlaps. Because as we move away from the sample the information in these overlaps decreases, the outer areas of the sample, where a large fraction of the beads sit, will be relatively worse stitched (as often we will need to rely on the default microscope displacements) than those near the sample, and therefore the former may dominate the registration error budget.

## Data availability

IMAXT aims to produce periodic data releases, including processed imaging data, segmentation catalogues, federated datasets, masks used for training segmentation neural networks, and so on. We also plan to open our infrastructure to external users to perform their analysis.

Information on these data releases and how to access the data can be accessed from https://imaxt.ast.cam.ac.uk/release.

## Code availability

All the code to perform stitching, registration and segmentation as well as additional tools is available in the IMAXT GitHub organization at https://github.com/IMAXT/under a GNU General Public License version 3 (GPLv3). In particular the STPT stitching and registration pipeline is available from https://github.com/IMAXT/stpt-mosaic-pipeline and the IMC nuclear channel segmentation pipeline in https://github.com/IMAXT/imc-segmentation-pipeline.

Additional information on our infrastructure, software tools, data model and applications is available from https://imaxt.github.io.

## Acknowledgements

I.V.G. is supported by a fellowship from the Ovarian Cancer Research Alliance. S.P.S. acknowledges support from the MSK Cancer Center Support Grant/Core Grant (P30 CA008748). N.A.W. acknowledges support from the UK Space Agency (ST/R004838/1) and the Science and Technologies Facility Council-UK Research and Innovation (ST/T003081/1).

This work was supported by Cancer Research UK [A24042, A21143]. G.J.H. is a Royal Society Wolfson Research Professor.

## Author information

### Consortia

#### CRUK IMAXT Grand Challenge Consortium

H. Raza Ali, M. Al Sa’d, S. Alon, Samuel Aparicio, G. Battistoni, S. Balasubramanian, R. Becker, Bernd Bodenmiller, E. S. Boyden, D. Bressan, A. Bruna, B. Marcel, Carlos Caldas, M. Callari, I. G. Cannell, H. Casbolt, N. Chornay, Y. Cui, A. Dariush, K. Dinh, A. Emenari, Y. Eyal-Lubling, J. Fan, E. Fisher, E. A. González-Solares, C. González-Fernández, D. Goodwin, W. Greenwood, F. Grimaldi, G. J. Hannon, O. Harris, S. Harris, C. Jauset, J. A. Joyce, E. D. Karagiannis, T. Kovačević, L. Kuett, R. Kunes, A. Küpcü Yoldaş, D. Lai, E. Laks, H. Lee, M. Lee, G. Lerda, Y. Li, J. C. Marioni, A. McPherson, N. Millar, C. M. Mulvey, F. Nugent, C. H. O’Flanagan, M. Paez-Ribes, I. Pearsall, F. Qosaj, A. J. Roth, Oscar M. Rueda, T. Ruiz, K. Sawicka, L. A. Sepúlveda, S. P. Shah, A. Shea, A. Sinha, A. Smith, S. Tavaré, S. Tietscher, I. Vázquez-García, S. L. Vogl, N. A. Walton, A. T. Wassie, S. S. Watson, T. Whitmarsh, S. A. Wild, E. Williams, J. Windhager, C. Xia, P. Zheng & X. Zhuang

## Contributions

E.A.G.S., A.D., C.G.F., A.K.Y., M.A.S, T.W. and N.A.W. built the software infrastructure, defined the data model and data flow procedures, wrote the data analysis pipelines and wrote the manuscript; N.M. installs and maintains the software and hardware infrastructure; STPT and IMC experiments were performed by A.F., C.M., M.P.R. and F.Q.; L.K. and J.W. contributed to the IMC data acquisition and analysis; I.F., D.G., A.R., I.V.G and S.W. contributed to the data analysis and infrastructure working group; E.A.G.S., D.B. and N.A.W defined the analysis system architecture, S.A., B.B., E.B, C.C., O.H., S.P.S, S.T., N.A.W. and G.H. directed the IMAXT project; and all authors read and approved the final manuscript.

## Ethics declarations

### Competing interests

S.P.S. is a founder and shareholder of Canexia Health Inc. No other author has competing interests to declare.

**Figure A1:**
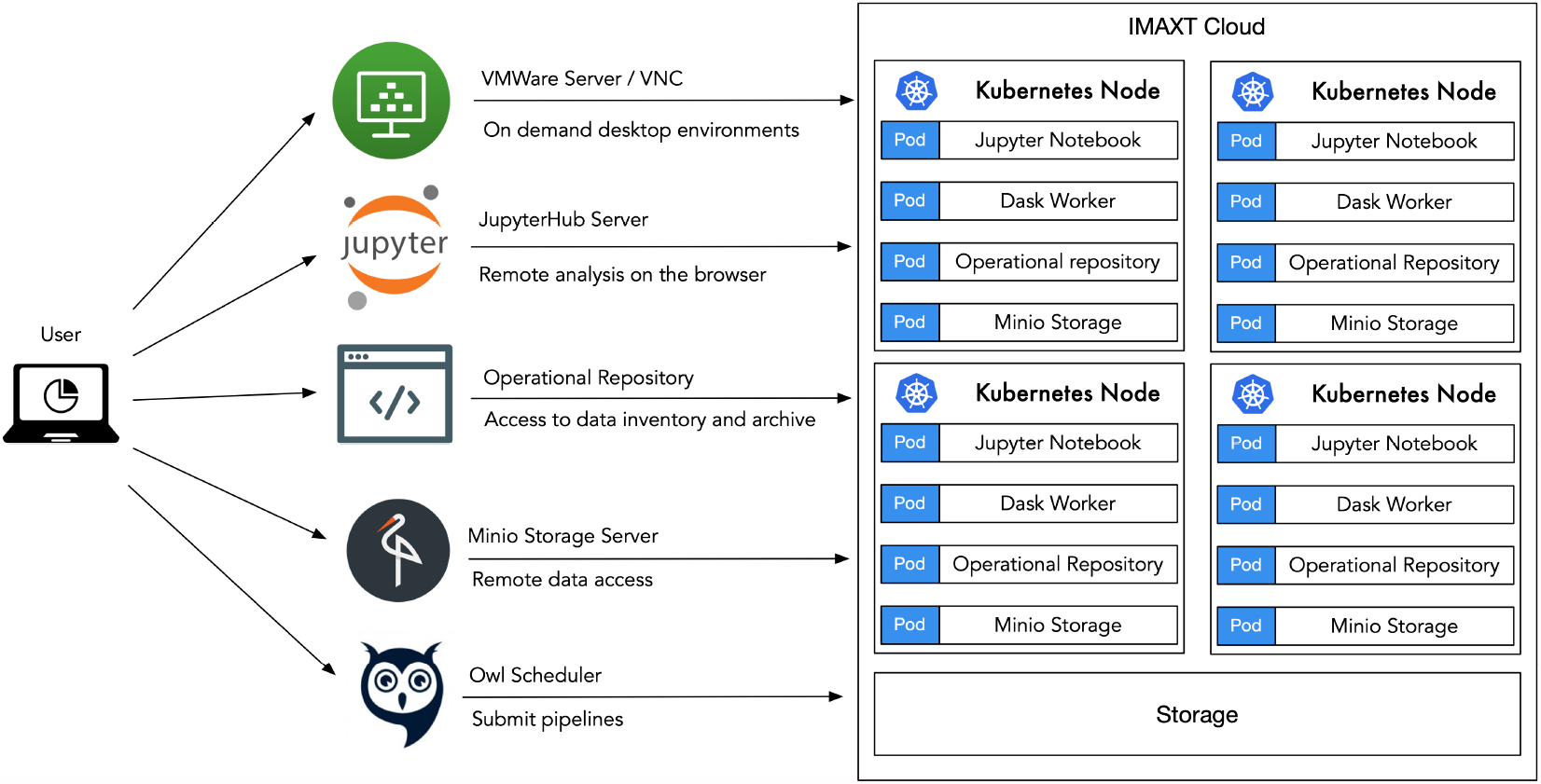
Services available to the user in the IMAXT Cloud. These include on demand remote desktops, Jupyter notebooks, job scheduler, archive and storage.

## Supplementary information

### A cloud for scientific exploitation

As figure A1 shows the IMAXT Cloud enables a range of services for the users. On demand Windows and Linux desktops environments with configurable resources allow to work close to where the data are with many software packages already installed and with a familiar look and feel. Jupyter notebook servers allow remote analysis and visualization in the browser. The operational repository and the associated database stores the metadata associated with the available datasets as well as results from the pipelines and allows for customized queries. We also provide utilities for transferring data and a custom job scheduler for submitting jobs to take full advantage of the computer power.

All the above runs in our own cluster based on Kubernetes that provides dynamic resource allocation based on the user and workload needs.

### Custom job scheduler

A job scheduler is an application that takes care of running unattended jobs in a compute cluster. Typically the scheduler will add jobs to a queue and run them when resources become available. Some widely used schedulers (mainly in HPC context) are Portable Batch System (PBS), Slurm Workload Manager, Condor, Moab, and many others.

In the context of cloud systems and in particular Kubernetes there is not a wide range of applications to run user submitted batch jobs. For our use such an application should be:

**Figure A2:**
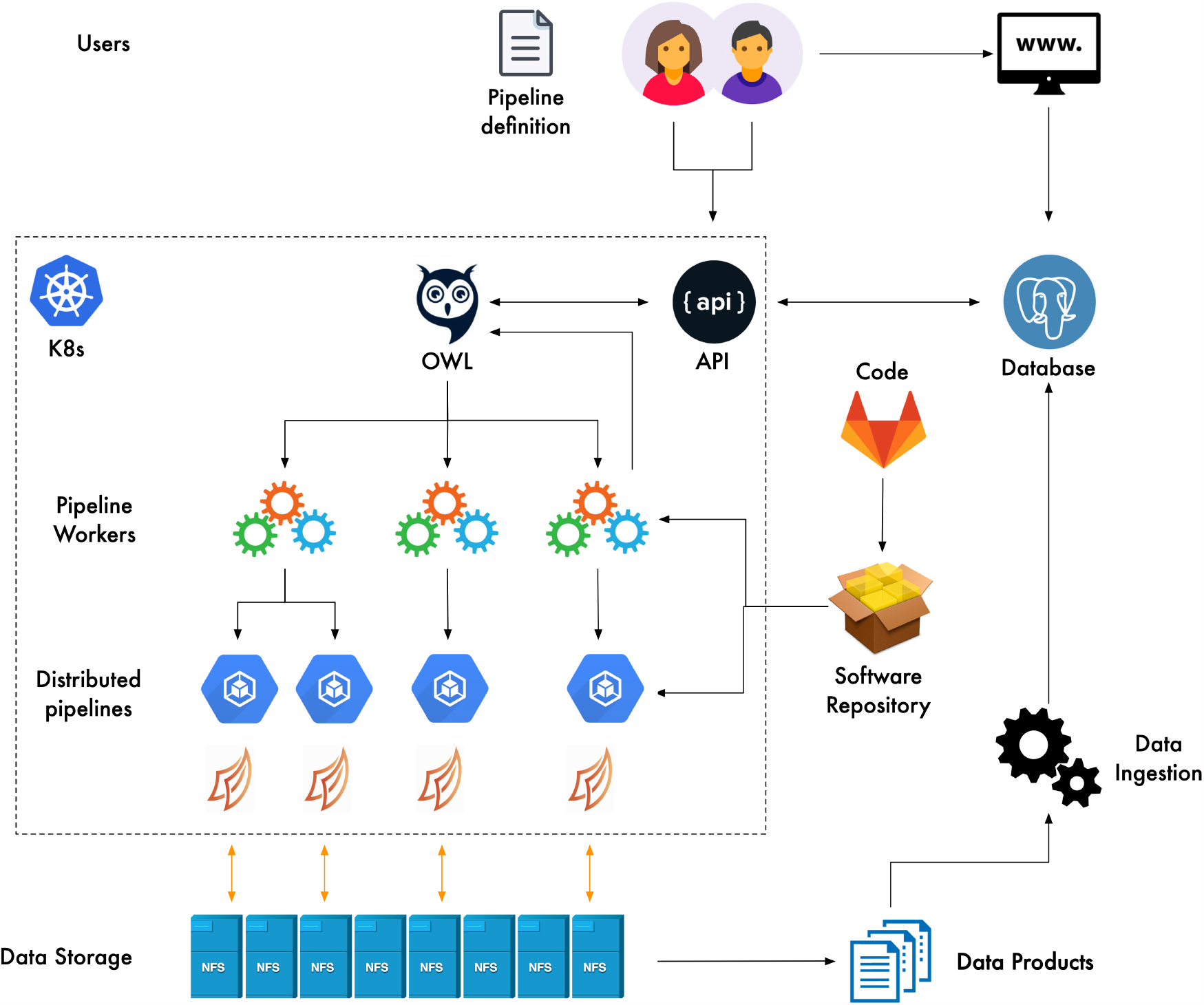
Pipeline scheduling. Users (or automatic processes) submit a Pipeline Definiton File (PDeF) that contains the name of the pipeline to submit, its arguments (location of input data, out-put, configurable parameters) and the cluster resources required. This is logged to a database that is queried by the scheduler (Owl) periodically. If there are available resources the pipeline is launched and the Dask workers are started. The progress is monitored and recorded in a database.

- Easy to use. Users should not be expected to know how the cluster is set up. A simple specification of how many CPUs and RAM should suffice in first instance with flexibility for more complex scheduling options.
- Remote access. Users should be able to submit, monitor and manage jobs from their own laptop or desktop without the need to login to a remote host.
- Dask support. It should support Dask jobs, but not exclusively.
- Integration with the rest of the system. The scheduler needs to create services and deployments in the K8s cluster, run jobs and interact with the archive authentication and the database.

For this purpose we have built the Owl job scheduler from scratch. Owl is a framework to execute jobs in a compute cluster backed up by Kubernetes and Dask. In essence Owl serves as very simple scheduler that accepts job descriptions, queues them and submits them for execution keeping track of progress.

In order to run a particular job (pipeline) users write a pipeline definition file, in summary, a file that specifies the pipeline to run and the arguments required for it to run as well as the compute resources required. This is submitted to the Owl API who logs the entry in a database and places the job in a queue. Owl checks periodically for pipelines in the queue and if they meet the scheduling requirements starts the allocation process. The main Owl process (a.k.a. Owl Scheduler) delegates the responsibility of running the pipeline to a pipeline worker Pod process that is in charge of contacting the K8s cluster, allocating the resources need and submitting the pipeline to the compute cluster.

The steps followed when running a pipeline are (see figure A2):

- The user submits a pipeline definition file (PDeF) in YAML format.
- A database is updated with the pipeline request. The Scheduler queries the database when slots are available and allocates pipelines.
- The Owl scheduler starts a pipeline worker in a Pod sending the pipeline definition file and extra configuration needed.
- The pipeline worker loads the pipeline code from a pip repository (either PyPi or our own private repository) and validates inputs against its schema.
- The Kubernetes cluster starts the requested number of Pod Dask workers.
- The pipeline worker starts a separate thread and runs the pipeline code. The code runs in the allocated containers using Dask taking advantage of parallelization over all workers.
- The main pipeline thread listens for heartbeat connections from the scheduler and waits for pipeline completion.
- The pipeline worker responds with the pipeline completion result to the scheduler.
- The scheduler stops the pipeline worker and logs an entry in the database.

Pipeline submission is done from the command line of any computer once the user is authenticated or from the Jupyter and involves writing the pipeline definition file that describes the job to run, the arguments required by the job and the resources requested. Job progress can be monitored either in the web or via the command line.

### Data model and data formats

Data obtained by the instruments are initially stored in their own local attached storage. They are then moved manually to a NAS storage system which is then copied manually to an upload folder. A source data uploader then transfers the data to our IoA image storage servers.

In Zarr datasets, the arrays are divided into chunks and compressed. These individual chunks can be stored as files on a filesystem, as objects in a cloud storage bucket or even in a database making it efficient for clusters of CPUs to access the data in parallel. The metadata are stored in lightweight .json files and allows all the metadata to be in a single location which requires just one read. Zarr works well on both local filesystems and cloud based object stores. Existing datasets can easily be converted to Zarr via Xarray’s Zarr functions. There are also existing functions to export Zarr to formats like HDF5 and we have written custom exporters to TIFF.

In summary the main advantages of Zarr are:

- Metadata is kept separate from data in a lightweight .json format
- Arrays are stored in a flexible chunked / compressed binary format
- Individual chunks can be retrieved independently in a thread safe manner
- The rate at which data can be extracted from a dataset scales linearly with the number of compute nodes which are reading from it simultaneously.

**Figure A3:**
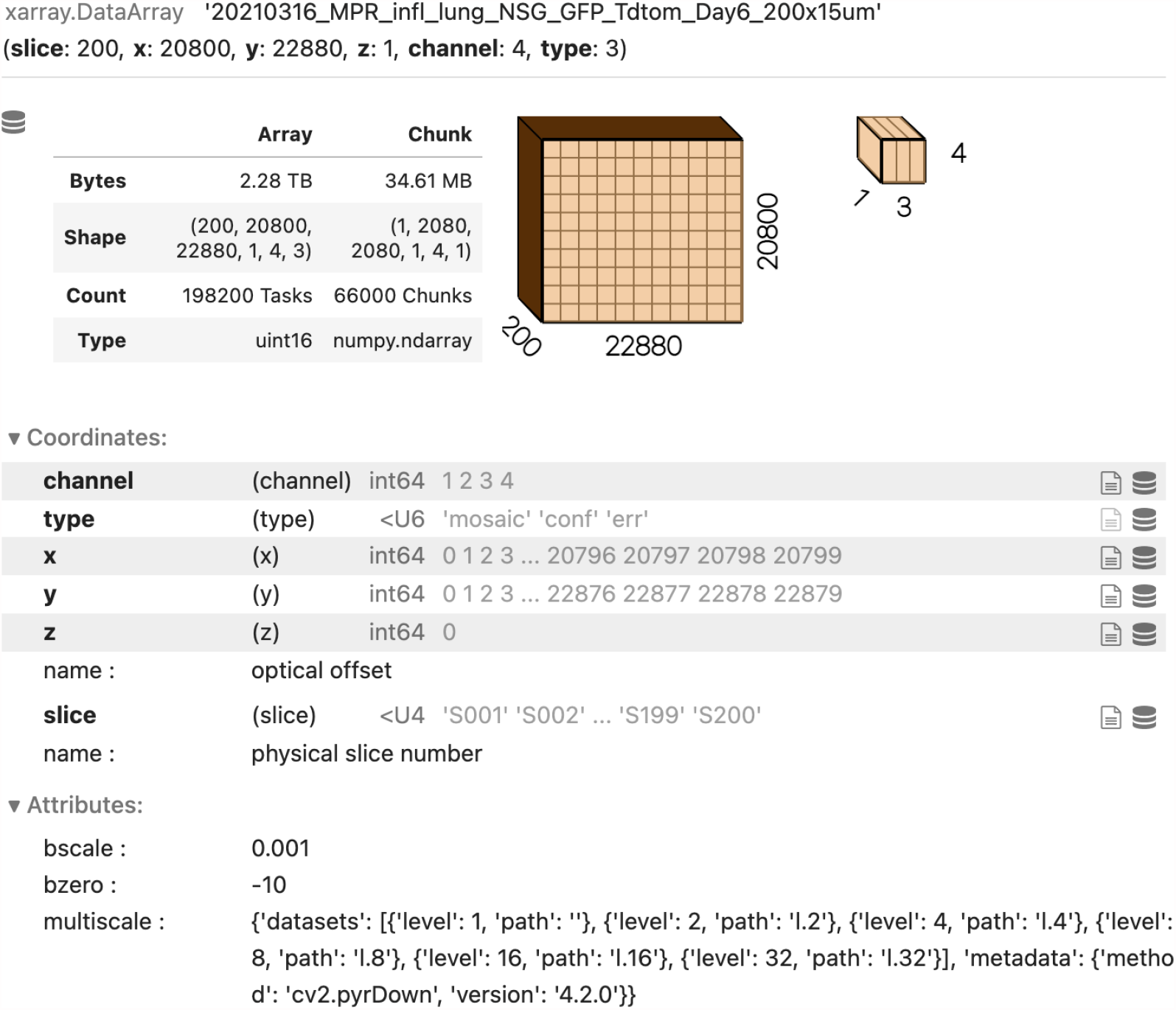
Representation of STPT dataset in Xarray. This particular STPT datasets contains 200 slices of 20800×22880 pixels and there are four channels. Together with the full image there is a confidence map and an error array associated of the same size. This dataset is divided into 66000 individual chunks that correspond to the same number of files on disk. Each chunk is a Dask array that is read from the storage only when necessary. This dataset amounts to a total of 2TB. Metadata are stored alongside the data.

Datasets are opened using the Xarray^30^ library in Python using the Zarr back-end. Xarray allows to store and read large chunked datasets in a cloud optimized fashion. Data are lazy-loaded, meaning that only the specific chunks of data are read when required and operations can be performed in paralell using Dask.

In order to further reduce the size of the datasets in disk we convert the data values to unsigned integers and record in the metadata the conversion values bscale and bzero so that float values can be recovered by multiplying the stored pixel value by bscale and adding bzero.

Additionally in order to support image visualization (e.g. using Napari, https://napari.org), we write additional data cubes downsampled a factor of 2^*n*^, *n* = 1, …, 5. These can be also exported to pyramidal TIFF.

## Footnotes

* Prior to IMC there is a high temperature drying process that we have found does not deform the sample slices appreciably

